# APOE genotype confers context-dependent neurovascular vulnerability in immune-vascularized human forebrain organoids

**DOI:** 10.1101/2025.05.08.652864

**Authors:** Haohui Fang, Xiangyu Liao, C Korin Bullen, Ruoyan Pu, Hu Wang, Juliana Condoleo, Sueanne Chear, Xueyi Chen, Yanjun Zhang, Sonya Zhang, Da Huo, Kadia Lissit, Alina Yang, Kateri Jarvis, Stewart Neifert, Yuejia Huang, William Bishai, Sanjay K. Jain, Ted M. Dawson, Valina L. Dawson, Jin-Chong Xu

**Author notes:** Correspondence should be addressed to: Neuroregeneration and Stem Cell Programs Institute for Cell Engineering Department of Neurology Johns Hopkins University School of Medicine 733 North Broadway, Suite 769 Baltimore, MD 21205. Co-first author (Haohui Fang, Xiangyu Liao, C Korin Bullen).

## Abstract

The APOE gene is a major genetic determinant of neurovascular and immune function, yet the mechanisms by which its isoforms modulate brain vulnerability to pathogenic stress remain incompletely understood. Here, we employ isogenic human iPSC-derived immune-vascularized—Forebrain Organoid-based Multicellular Assembled Cerebral Organoids (FORMA-COs)—to dissect isoform-specific responses to a clinically relevant viral challenge. We find that APOE2/2 and APOE4/4 FORMA-COs exhibit heightened viral RNA burden and distinct neuroinflammatory profiles compared to APOE3/3. Specifically, APOE4/4 promotes IL-1α and VEGFA induction, whereas APOE2/2 leads to elevated TNF-β and VEGFA protein accumulation, indicating divergent pathways of injury. Integrated transcriptomic analyses, combined with known and predicted APOE protein–protein interaction networks, reveal genotype-dependent enrichment of cytokine signaling, angiogenic remodeling, and immune dysregulation. In vivo validation using humanized mouse models corroborates APOE genotype– specific vascular remodeling, microglial activation, and oligodendrocyte perturbation. These findings demonstrate that APOE genotype confers context-specific susceptibility to neuroimmune and vascular injury, providing insight into genetic risk mechanisms underlying infection-related and neurodegenerative brain disorders.

## Introduction

Apolipoprotein E (APOE) is a multifunctional lipid transport protein with critical roles in the central nervous system (CNS), where it regulates cholesterol homeostasis, synaptic stability, immune responses, and blood-brain barrier (BBB) integrity. The three common human APOE isoforms—APOE2, APOE3, and APOE4— arise from coding polymorphisms in the APOE gene and exert distinct effects on cellular and tissue physiology. Among these, APOE4 is the strongest common genetic risk factor for late-onset Alzheimer’s disease (AD), while APOE2 is often regarded as protective^1–7^. However, emerging evidence challenges this binary classification, indicating that both APOE2 and APOE4 can confer increased vulnerability under specific pathological conditions, particularly those involving vascular compromise and immune activation^8–14^.

Recent studies have broadened the clinical relevance of APOE genotype beyond neurodegenerative disorders to include infectious and systemic diseases. In particular, a landmark study demonstrated that mice expressing human APOE2 or APOE4 exhibit increased mortality, higher viral burden, and impaired immune responses following SARS-CoV-2 (hereafter SCV2) infection compared to APOE3-expressing controls^15^. These findings, corroborated by population-level genetic data, suggest that APOE genotype modulates host responses to pathogenic insults in a manner that is highly context-dependent. Despite these advances, the molecular and cellular mechanisms underlying isoform-specific responses— especially within human neurovascular compartments—remain poorly understood.

To address this gap, we employed isogenic human induced pluripotent stem cell (iPSC)-derived immune-vascularized forebrain organoids—Forebrain Organoid-based Multicellular Assembled Cerebral Organoids (FORMA-COs)—to model how APOE isoforms influence neuroimmune and vascular responses to defined pathogenic stress. These assembloids recapitulate key components of the human neurovascular unit, incorporating neurons, astrocytes, microglia, endothelial cells, and pericytes. Using SCV2 as a clinically relevant and mechanistically well-characterized viral challenge, we compared APOE2/2, APOE3/3, and APOE4/4 FORMA-COs for viral RNA burden, cytokine and growth factor signaling, glial reactivity, and vascular remodeling. Findings were validated in an in vivo humanized ACE2 (hACE2) mouse model to assess translational relevance.

We show that APOE2 and APOE4 confer distinct but converging patterns of vulnerability, characterized by increased viral RNA accumulation, dysregulated cytokine responses, and aberrant VEGF signaling. While APOE4 organoids mounted exaggerated IL-1α and angiogenic responses, APOE2 organoids showed elevated TNF-β production and VEGFA protein accumulation. These divergent signatures were reflected in infected mouse brains, which exhibited APOE-linked fibronectin redistribution, microglial activation, and myelin-associated changes. Together, our results demonstrate that APOE genotype governs context-dependent neurovascular susceptibility to pathogenic stress, offering a mechanistic bridge between genetic risk and heterogeneous outcomes in infection and neurodegenerative disease.

## Results

### Generation of Immune-Vascularized Human Forebrain Organoids

To model human-specific neurovascular and immune responses in a 3D forebrain context, we developed a protocol for generating immune-vascularized multicellular organoids, termed FORMA-COs (Figure 1A). This platform was designed to recapitulate pathological features observed in SCV2–positive postmortem brain tissues, including microgliosis, astrogliosis, vascular injury, neuronal loss, and demyelination^16–36^ (Figure S1A).

**Figure 1.**
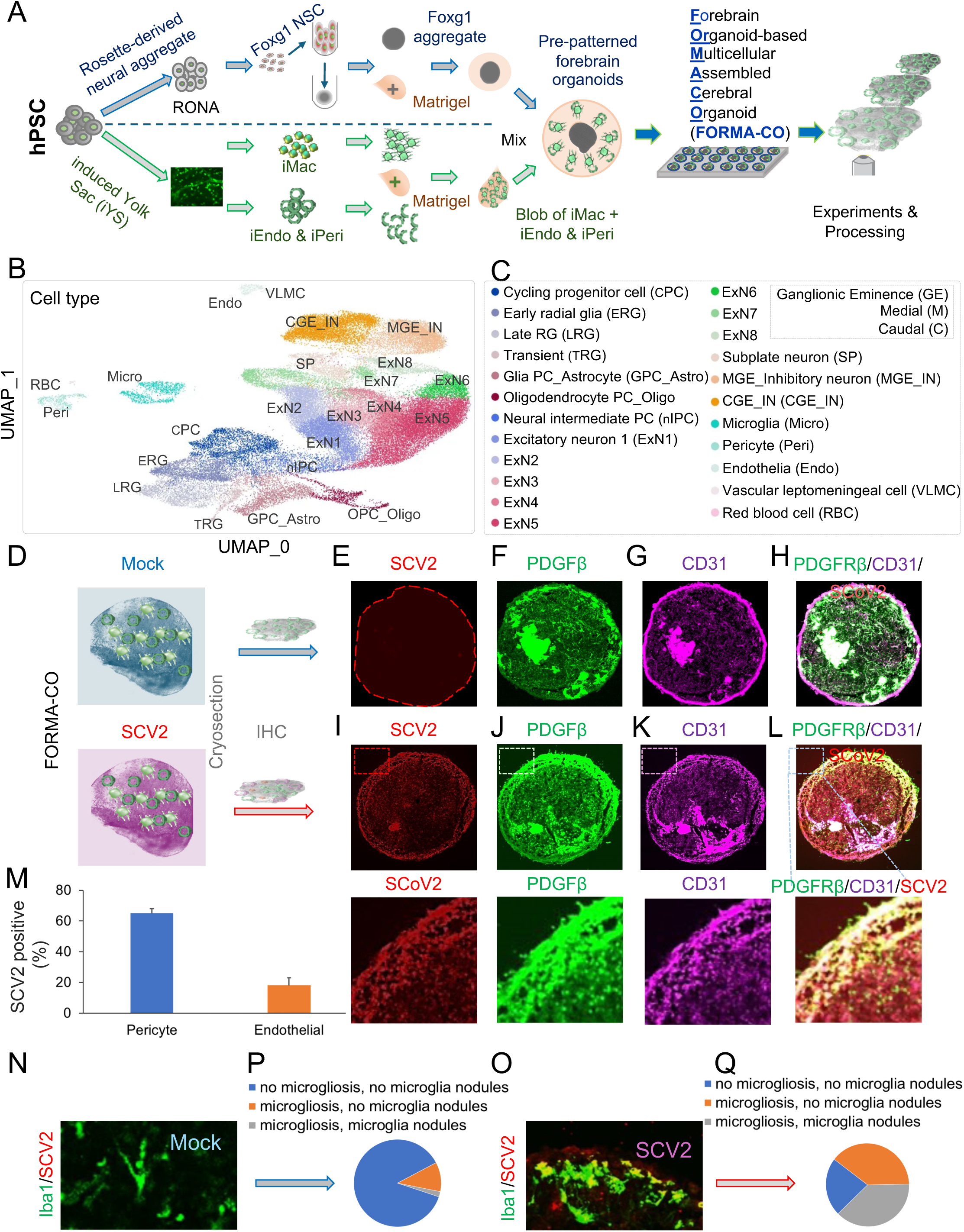
**Generation and SCoV2 infection of immune-vascularized forebrain organoids (FORMA-COs).** (A) Schematic of FORMA-CO assembly combining FOXG1⁺ neural aggregates with iPSC-derived macrophages (iMac), endothelial cells (iEndo), and pericytes (iPeri). (B–C) UMAP plots showing diverse neurovascular and immune cell populations. (D) Workflow for SCoV2 versus mock infection. (E–L) Immunofluorescence shows SCoV2 localization with pericyte (PDGFRβ) and endothelial (CD31) markers. (M) Quantification of SCoV2⁺ vascular cells. (N) Colocalization of Iba1⁺ microglia with virus; pie charts summarize microgliosis.

FORMA-COs were derived through the directed differentiation of human induced pluripotent stem cells (iPSCs) into multipotent forebrain neural stem and neural progenitor cells expressing FOXG1 and Nestin (Figure S1B). These patterned progenitors retained multipotency and gave rise to forebrain neurons^37^, GFAP⁺ astrocytes, and OLIG2⁺oligodendrocyte-lineage cells during terminal differentiation (Figure S1C–D).

In parallel, we induced yolk sac–like tissues from iPSCs to generate mesodermal derivatives resembling early extraembryonic hematovascular lineages. These iPSC-derived yolk sac (iYS) included CD31⁺ endothelial cells and CD45⁺ hematopoietic progenitors, which further differentiated into supernatant macrophage-like cells (iMac), as well as additional endothelial (iEndo) and pericytic (iPeri) cells localized along the basement membrane within the same culture system (Figures S1E–G). Transcriptomic comparisons via principal component analysis revealed strong concordance between our iYS tissues and primary human yolk sac samples (Figure S1F). Moreover, RNA sequencing of iMacs confirmed high expression of canonical microglial genes^38^, validating their myeloid lineage identity (Figure S1G).

To reconstruct the immune-vascular interface, these components were temporally integrated into developing forebrain organoids. Pre-patterned FOXG1-NPC derived forebrain organoids were embedded in Matrigel supplemented with iMacs, iEndo, and iPeri to facilitate coordinated cellular integration and structural maturation. To monitor integration dynamics, we introduced fluorescently labeled iMacs (RFP⁺) and iEndo/iPeri (GFP⁺) and tracked their localization over time. Imaging revealed broad incorporation of vascular and immune cells throughout the organoid parenchyma, with microglia clustering preferentially along vascular surfaces (Figure S1H–K). Quantification of endothelial-associated microglia (EAMs) over time supported a model of immune–vascular co-migration during organoid maturation (Figure S1L). A schematic summary illustrates the modular assembly strategy and temporal integration steps of the FORMA platform (Figure S1M).

To assess the cellular composition of mature FORMA-COs, we performed deconvolution of bulk RNA-seq using a single-cell reference from the developing human cortex. This analysis revealed a complex and reproducible cell-type landscape (Figure 1B–C), including forebrain neurons and glia, endothelial cells, pericytes, and macrophage-lineage immune cells. Dimensionality reduction via UMAP and marker-based annotation confirmed successful co-differentiation and spatial assembly of these diverse populations.

Collectively, these data demonstrate that FORMA-COs constitute a robust and tractable platform for modeling integrated brain-intrinsic and neurovascular-immune interactions in a human-specific, physiologically relevant 3D system.

### Temporal Dynamics of SCoV2 Replication in FORMA-COs

We next profiled viral replication kinetics in FORMA-COs following SCoV2 exposure. RT-qPCR revealed a rapid rise in viral RNA at 6 hours post-infection, followed by a steep decline by day 3 and undetectable levels by day 6 (Figure S1N). Consistent with this, TCID₅₀-based titration showed peak infectious viral load at 6 hours with rapid attenuation thereafter (Figure S1O). These results suggest that FORMA-COs support early stages of viral replication, potentially followed by intrinsic antiviral responses that limit spread.

### FORMA-COs as a Platform to Model SCV2 Neuroinvasion

To assess the utility of FORMA-COs in modeling viral neuropathogenesis, we exposed immune-vascularized organoids to SCV2 and conducted multiplex immunohistochemical analyses to localize infection and characterize host responses (Figure 1D). Infected organoids exhibited prominent viral signal within pericyte (PDGFRβ⁺) and endothelial (CD31⁺) compartments (Figure 1E–L). Viral particles were frequently detected within or adjacent to vascular-associated cells, and quantitative analysis confirmed infection of both pericyte and endothelial populations (Figure 1M), implicating these components as early cellular targets of SCoV2 within the neurovascular unit.

In addition to vascular involvement, SCoV2 exposure induced marked microglial activation. Iba1⁺ microglia displayed reactive morphologies and aggregated into nodular clusters in infected FORMA-COs, a phenotype absent in mock-treated controls (Figure 1N). These observations mirror reported patterns of microgliosis in postmortem COVID-19 brain tissues and support the relevance of FORMA-COs for recapitulating innate immune responses to viral invasion.

Finally, confocal imaging revealed neuron-specific infection with SCV2 signal colocalizing with β-tubulin and nuclear stain TO-PRO-3, albeit at low or near-undetectable frequencies (Figure S1P). Three-dimensional reconstructions confirmed spatial viral localization within neuronal somas and processes, implicating direct neurotropism within the organoid model.

Together, these findings demonstrate that FORMA-COs provide a highly adaptable and human-specific platform to model neurotropic viral infections. The observed cellular and pathological features benchmark closely with those in SCV2-positive postmortem brain tissues from individuals with confirmed COVID-19, enabling mechanistic investigation of early immune-vascular injury within the developing human forebrain.

### Time-Resolved Transcriptional Profiling Reveals Multisystem Dysregulation Following SCV2 Infection in FORMA-COs

To uncover the molecular mechanisms underlying SCV2–induced neurovascular pathology, we performed a time-resolved gene expression analysis using a customized NanoString panel targeting neural, immune, and vascular response genes in FORMA-COs. Total RNA was extracted from SCV2–infected and mock-infected organoids at 6-, 24-, and 72-hours post-infection (HPI), followed by hybridization to multiplexed barcoded probes and digital transcript quantification (Figure 2A).

**Figure 2.**
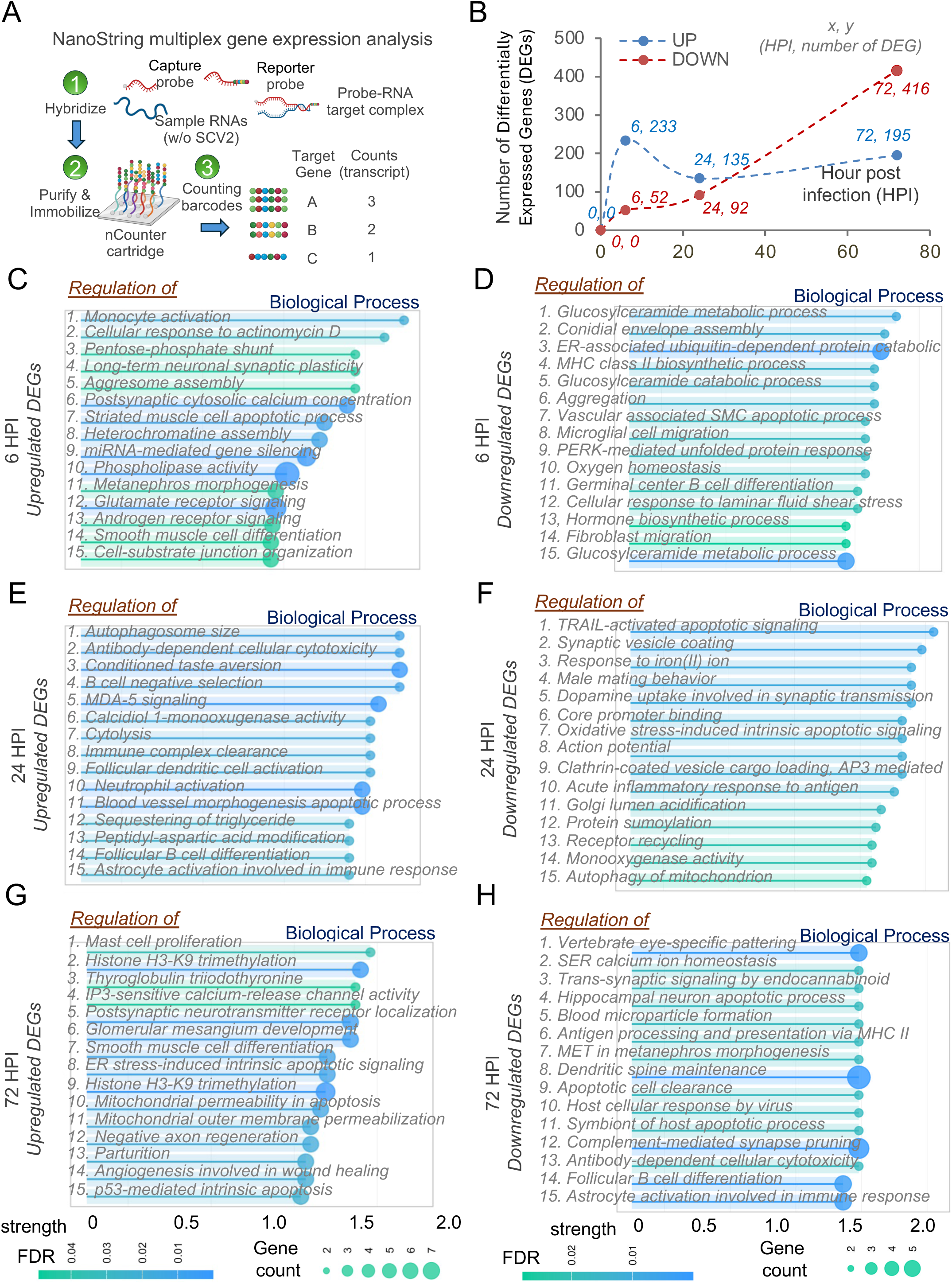
**Time-resolved transcriptional responses to SCoV2 infection in FORMA cerebral organoids.** (A) NanoString assay workflow for profiling immune and neuronal gene expression. (B) DEG dynamics across 6, 24, and 72 hours post-infection (HPI); peak response at 72 HPI. (C–H) GO enrichment of DEGs shows time-dependent activation of immune, neuronal, vascular, and apoptotic pathways. Upregulated genes reflect inflammation and tissue injury; downregulated genes involve synaptic signaling and metabolic regulation.

The transcriptional response exhibited a clear temporal progression, with the number of differentially expressed genes (DEGs) increasing over time. By 72 HPI, we observed peak molecular perturbation, with 416 genes significantly downregulated and 195 upregulated (Figure 2B). These dynamics suggest that SCV2 elicits a delayed but intensifying cellular response in the neural tissue environment^39–41^.

Gene ontology (GO) enrichment analysis of DEGs revealed distinct biological processes at each time point. At 6 HPI, upregulated genes were enriched for innate immune activation—including monocyte recruitment^42–44^ and interferon signaling^45, 46^—alongside chromatin remodeling and miRNA regulation, consistent with early hyperinflammatory responses (Figure 2C). In contrast, downregulated genes indicated suppression of MHC class II antigen presentation, B cell differentiation, and endoplasmic reticulum (ER) stress responses, as well as early signs of vascular injury and neuronal dysfunction^47, 48^ (Figure 2D). These findings suggest that SCV2 rapidly disrupts immune homeostasis, neurovascular stability, and intracellular stress regulation^49–51^.

By 24 HPI, immune dysregulation intensified. Upregulated pathways reflected enhanced antibody-dependent cytotoxicity, astrocyte-mediated immune signaling, and compromise of vascular integrity (Figure 2E). Concurrent downregulation affected pathways related to synaptic signaling, dopamine neurotransmission, mitochondrial homeostasis, protein trafficking, and endocrine regulation (Figure 2F), indicating growing systemic involvement across neural and metabolic networks.

At 72 HPI, transcriptional signatures revealed widespread tissue remodeling, systemic immune infiltration, and sustained neuronal injury. Upregulated genes were enriched for mast cell proliferation, histone modifications, ER stress responses, and chromatin remodeling (Figure 2G), indicating escalating immune and epigenetic disruption. Downregulated genes pointed to deficits in synaptic maintenance, dendritic spine structure, hippocampal function, and vascular remodeling (Figure 2H), suggesting convergence of neuroinflammation, neurodegeneration, and cerebrovascular instability.

Together, these results define a temporally structured and progressively worsening transcriptional landscape in SCV2-infected FORMA-COs—one that bridges innate and adaptive immune activation with microvascular damage and neurofunctional decline.

### Pathway-Level Dissection of SCV2 Responses in FORMA-COs Reveals Oxidative Stress, Neurodegeneration, and Vascular Remodeling

To further dissect the systemic responses driving SCV2-induced pathology in FORMA-COs, we performed GO and Reactome pathway enrichment analyses on the time-resolved NanoString dataset (Figure S2). This approach enabled identification of temporally coordinated pathway activation across immune, neural, and vascular compartments.

Upregulated DEGs consistently showed enrichment for immune activation, cytokine production, and B cell–mediated responses across all time points (Figure S2A). Early disruption of neural and glial developmental pathways—including those involved in oligodendrocyte maturation and neural progenitor proliferation—was evident at 6 HPI, suggesting potential long-term neurodevelopmental consequences. As infection progressed, transcriptional shifts reflected increasing engagement of adaptive immunity, gliogenesis, and tissue remodeling by 72 HPI.

Downregulated DEGs followed similarly structured temporal patterns (Figure S2B). At 6 HPI, downregulated pathways were dominated by oxidative stress regulation, reactive oxygen species (ROS) metabolism, and neuronal apoptosis. These signatures persisted and intensified at 24 HPI, alongside increased engagement of lysosomal degradation and metabolic stress pathways—indicative of progressive mitochondrial dysfunction and cumulative neurotoxicity.

Reactome analysis of upregulated DEGs revealed early activation of interleukin signaling, MAPK cascades, and platelet activation at 6 HPI (Figure S2C). By 24 HPI, persistent NF-κB signaling and chromatin modification became dominant, suggesting a transition from acute proinflammatory signaling to epigenetic reprogramming and cellular stress adaptation.

Conversely, downregulated Reactome pathways (Figure S2D) showed early suppression of cytokine, chemokine, and interferon signaling—despite concurrent immune activation— suggesting feedback disruption or viral immune evasion. By 24 HPI, this suppression extended to core transcriptional machinery, including RNA polymerase II–mediated transcription, SUMOylation, and mRNA processing, indicating widespread transcriptional and post-transcriptional dysregulation.

Temporal mapping of select Reactome terms provided further resolution of the evolving molecular landscape. At 6 HPI, antiviral and apoptotic pathways such as IFIH1/DDX58-mediated interferon signaling and mitochondrial caspase-8 activation were prominently upregulated, while anti-inflammatory mediators (e.g., CD163) and glycosphingolipid metabolism were suppressed (Figure S2E, H). By 24 HPI, interferon signaling persisted, accompanied by intensified repression of mRNA splicing, SUMOylation, and transcriptional elongation machinery, indicating increasing disruption of post-transcriptional regulation (Figure S2F, I). At 72 HPI, upregulated genes pointed to impairment of NMDA receptor function, gap junction communication, and synaptic plasticity, alongside activation of DNA repair and purinergic signaling pathways (Figure S2G). Concurrently, downregulated pathways included NFE2L2/HMOX1-mediated oxidative stress response, interferon signaling, insulin regulation, and cytoskeletal remodeling (Figure S2J), highlighting a convergence of neuroimmune dysfunction, oxidative damage, and metabolic dysregulation in the later phase of SCV2 infection.

Together, these data demonstrate that SCV2 infection triggers a temporally structured but pathologically escalating impact across neural, immune, and vascular domains. FORMA-COs faithfully capture these multifaceted dynamics, providing a powerful platform to dissect early antiviral defense, delayed neurovascular injury, and persistent transcriptional dysfunction— hallmarks of COVID-19–associated neuropathology [citations].

### Network-Based Analysis Reveals APOE-Centered Gene Modules Linking Neuroimmune and Vascular Pathologies

To identify key regulatory hubs and biological processes disrupted by SCV2 infection within the immune-vascularized brain organoids, we conducted a network-based gene enrichment analysis incorporating betweenness centrality, functional clustering, and pathway enrichment. Using integrated transcriptomic data from FORMA-COs, we constructed gene interaction networks and calculated betweenness centrality to prioritize influential molecular nodes. Betweenness centrality analysis identified CTNNB1, TNF, and HIF1A as the top three central regulators, reflecting their critical roles in bridging diverse biological modules (Figure 3A). Additional high-centrality nodes included TLR4, APOE, FYN, PTPRC, CD8A, RAB7A, and LAMP1, highlighting the convergence of inflammatory, immune, and vascular signaling pathways. Notably, APOE emerged as a prominent hub, suggesting a context-dependent regulatory function during neurovascular inflammation—a finding aligned with population-level genetic studies implicating APOE genotype in modulating host susceptibility to pathogenic stressors.

**Figure 3.**
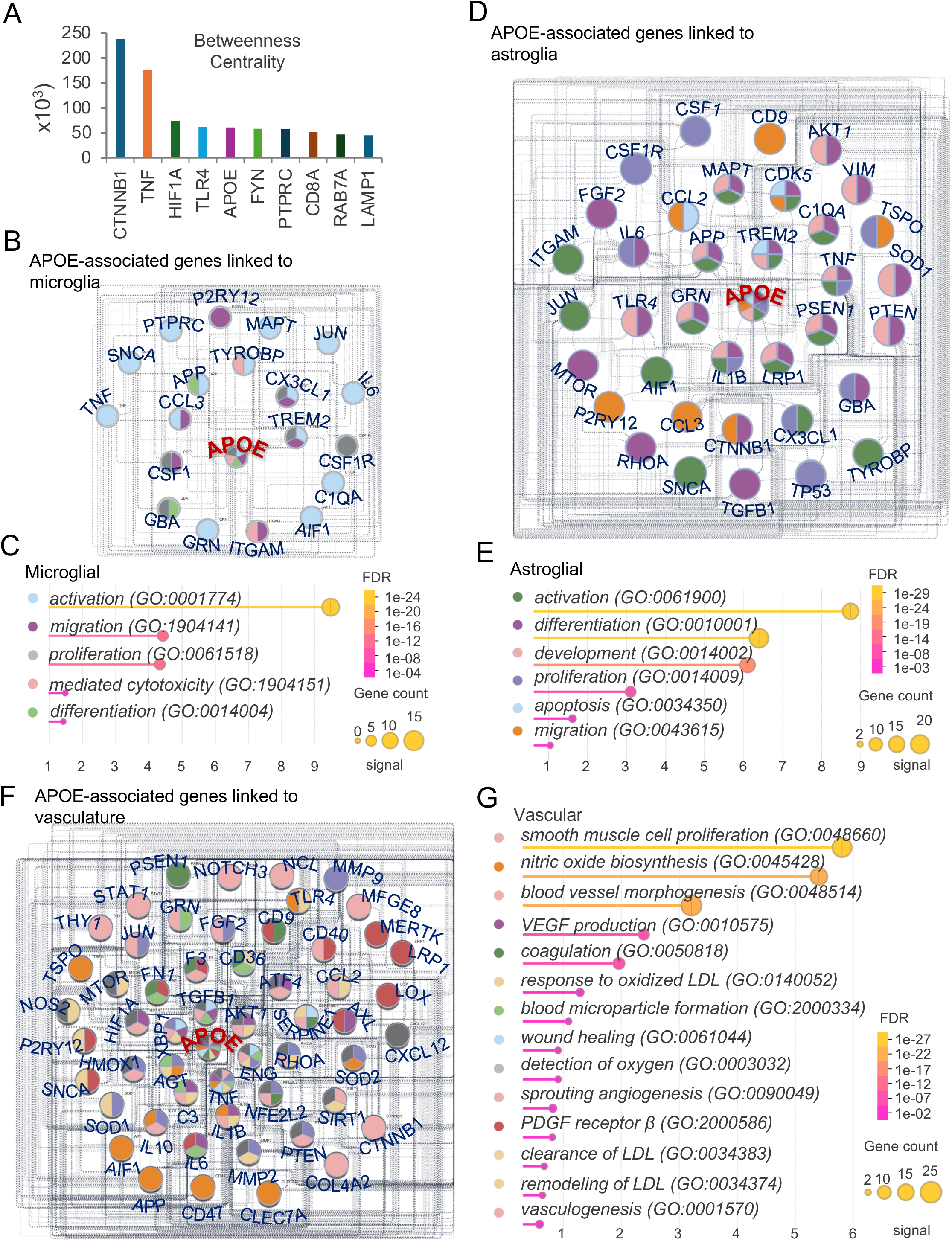
**Network analysis of APOE-centered responses to infection.** (A) Betweenness centrality highlights CTNNB1, TNF, and HIF1A as key regulators. (B–C) APOE-centered microglial network shows links to inflammation, phagocytosis, and neurodegeneration. (D–E) Expanded APOE network includes astrocytic and immune-related hubs (e.g., IL6, MAPT, TREM2) with enriched glial and cytokine pathways. (F–G) Enrichment of vascular-related GO terms implicates APOE in angiogenesis, endothelial signaling, and vascular remodeling.

Focusing on microglial gene networks, we constructed an APOE-centered interactome that revealed tight connectivity with genes involved in inflammation, phagocytosis, and neurodegeneration (Figure 3B). Interacting partners included canonical regulators such as TREM2, APP, MAPT, GRN, IL6, and CX3CL1, all of which are implicated in neurodegenerative disease pathophysiology [citation]. GO enrichment analysis of this APOE-centered microglial module revealed significant overrepresentation of biological processes related to microglial activation, migration, proliferation, differentiation, and cytotoxicity (Figure 3C), suggesting APOE’s potential role in coordinating both protective and pathogenic microglial functions.

Expanding the analysis beyond microglia, we observed that the broader APOE-centered network incorporated genes involved in astrocytic, endothelial, and neuronal signaling (Figure 3D). GO enrichment of this expanded network revealed strong representation of astroglial activation, proliferative signaling, and apoptotic regulation (Figure 3E), pointing to a wider modulatory role for APOE across multiple glial lineages under inflammatory conditions.

Importantly, vascular-related pathways emerged as prominent features within the APOE interactome. Enrichment analyses identified biological processes including endothelial signaling, extracellular matrix remodeling, oxidative stress response, angiogenesis, and inflammatory vascular remodeling (Figure 3F–G).

Key contributing genes included HIF1A, FGF2, MMP9, NOS2, STAT1, and TGFB1, supporting a role for APOE not only in neuroimmune regulation but also in the maintenance—and potential disruption—of cerebrovascular homeostasis.

Collectively, these network-based analyses position APOE as a central integrator of immune activation, neurodegeneration, and vascular dysfunction in response to SCV2 infection. These findings expand the traditional view of APOE’s role in Alzheimer’s disease, establishing a broader framework for its function in neuroimmune and neurovascular regulation.

### Functional Segregation of APOE Interactomes Highlights Distinct Neuroimmune Modules

To further delineate the functional breadth of APOE’s interaction partners, we constructed a series of specialized APOE-centered networks enriched for neuronal, glial, and immune-specific processes (Figure S3). This modular segregation enabled visualization of how APOE coordinates parallel biological systems in response to pathogenic stimuli.

The neuronal/oligodendroglial APOE-centered network revealed strong connectivity to genes involved in synaptic integrity, neurotransmission, axon-glia interactions, and myelination (Figure S3A). Key nodes included DLG4, GRIN2B, PRNP, and NEFL, underscoring APOE’s role in maintaining neuronal excitability and structural plasticity. Additional connections to UBQLN1 and MSR1 implicated APOE in axonal maintenance and myelin stability. GO enrichment analysis highlighted pathways such as synaptic pruning, NMDA receptor regulation, and amyloid-beta clearance—signatures often linked to neurodegenerative vulnerability.

In contrast, a leukocyte-centered APOE network (Figure S3B) captured genes governing immune cell activation, chemotaxis, and effector responses. Genes such as PTPRC, CCL5, CX3CL1, and IL6 showed strong immune relevance, consistent with evidence of immune dysregulation in both COVID-19 and APOE-associated neuroinflammatory conditions. A dedicated neutrophil/eosinophil module (Figure S3C) identified granulocyte-mediated processes, with genes like CCL2, CSF1, CX3CL1, and TNF contributing to migration, adhesion, and cytokine production. Similarly, a monocyte-centered network (Figure S3D) highlighted innate immune mechanisms such as chemotaxis, phagocytosis, and inflammatory signaling, involving nodes like AIF1, CSF1R, and CLEC7A. The macrophage-specific network (Figure S3E) reinforced APOE’s role in regulating myeloid cell function, linking to genes including TREM2, CD36, and GRN. GO analysis indicated enrichment for macrophage activation, foam cell differentiation, and pro-inflammatory cytokine secretion. Within the adaptive immune compartment, the B/T cell APOE network (Figure S3F) connected APOE to genes regulating lymphocyte activation and cytokine output, including CD4, FOXP3, and PTEN, suggesting potential cross-talk between innate and adaptive immune responses in neuroinflammatory contexts. Finally, the cytokine signaling module (Figure S3G) underscored APOE’s involvement in regulating inflammatory cascades. Enriched pathways included IFN-γ, IL-6, IL-1β, IL-17, and IL-23 signaling, with central hubs such as IL1B, STAT1, and TLR4 reflecting coordinated dysregulation of both innate and adaptive immunity during infection.

Together, these functionally segregated APOE-centered interactomes highlight the gene’s remarkable versatility as a molecular integrator across neuronal, glial, and immune networks. This network plasticity likely underlies APOE’s context-dependent roles in aging, neurodegeneration, and susceptibility to infectious insults such as SCV2 [citations].

### APOE Genotype Influences SCV2 RNA Accumulation, Host Gene Expression, and Neuroimmune Signaling in FORMA-COs

To determine how APOE genotype modulates neuroimmune responses to viral infection, we utilized an isogenic human iPSC-derived FORMA-CO platform incorporating homozygous APOE2/2, APOE3/3, and APOE4/4 genotypes (Figure 4A). This approach enabled genotype-controlled comparisons across neural, immune, and vascular cell types under SCV2 challenge. APOE allele identity was confirmed by Sanger sequencing at codons 112 and 158 (Figure S4A), corresponding to SNPs rs429358 and rs7412, which define the canonical APOE isoforms.

**Figure 4.**
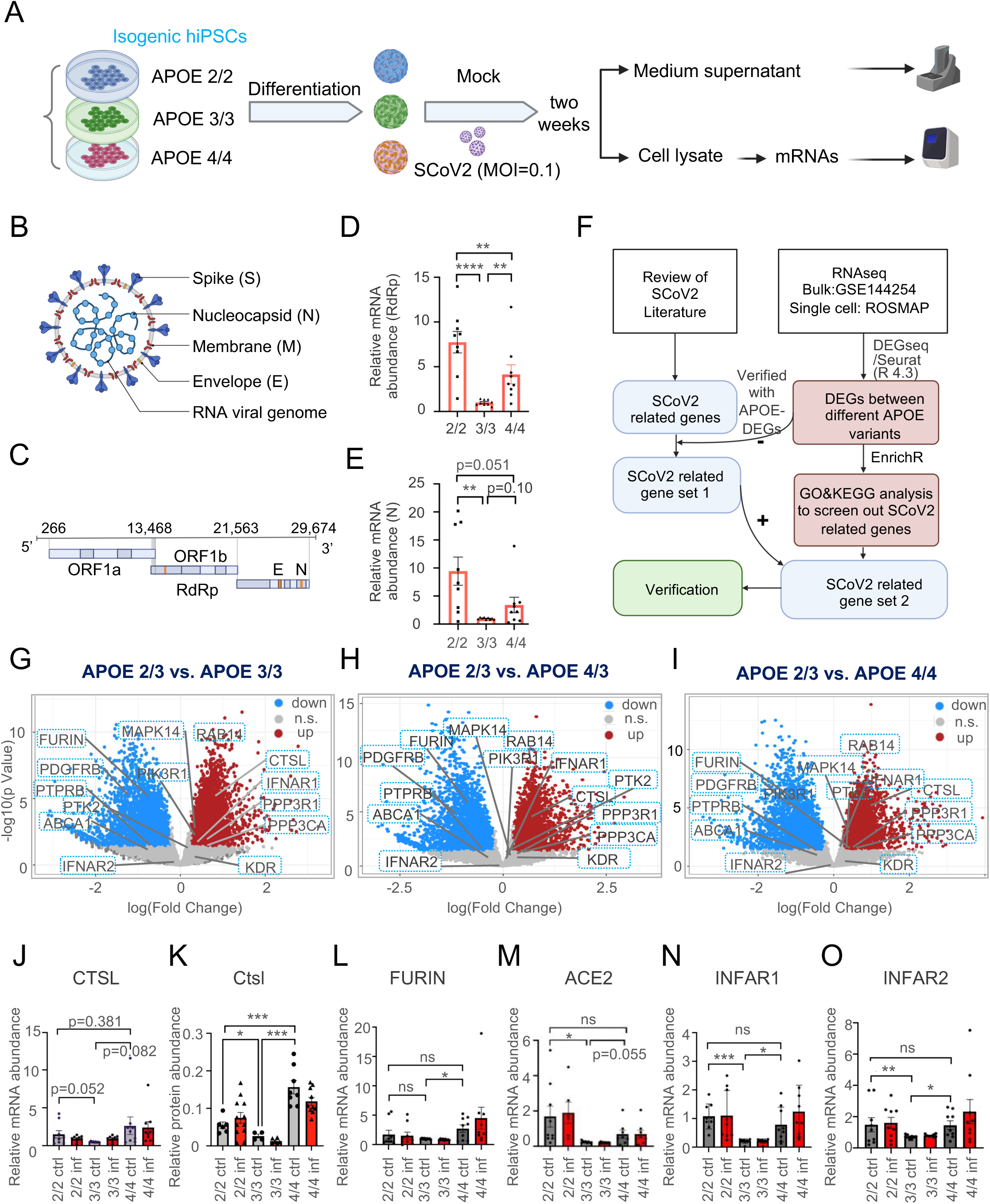
**APOE Genotype Modulates Viral RNA Accumulation and Host Responses in Isogenic FORMA-COs.** (A) Schematic of isogenic APOE2/2, 3/3, and 4/4 iPSC-derived FORMA-CO pipeline. (B-C) Diagram of SARS-CoV-2 structure and genome organization. (D-E) APOE2/2 and 4/4 FORMA-COs show elevated SARS-CoV-2 Rdrp and N RNA levels vs. APOE3/3. (F) Workflow for identifying SARS-CoV-2–responsive genes influenced by APOE genotype. (G-I) Volcano plots show DEGs in APOE2/3 vs. 3/3, APOE2/3 vs. 4/4, and APOE2/3 vs. 4/4 comparisons, revealing immune and vascular gene differences. (J-K) CTSL mRNA and protein levels trend higher in APOE4/4 and APOE2/2 controls, but infection does not alter expression. (L) FURIN mRNA is upregulated in APOE4/4 controls vs. APOE3/3. Infection has minimal additional effect. (M) ACE2 mRNA is elevated in APOE2/2 controls and trends higher in APOE4/4 vs. 3/3. (N-O) IFNAR1 and IFNAR2 mRNA show significantly lower expression in APOE3/3 than in APOE2/2 and 4/4; infection does not alter levels.

We first assessed whether APOE genotype influenced viral load by quantifying SCV2 RNA in infected FORMA-COs. SCV2 contains essential genomic elements for replication, including RdRp and N genes (Figure 4B–C). Quantitative analysis revealed significantly higher RdRp RNA levels in both APOE2/2 and APOE4/4 organoids relative to APOE3/3, with APOE2/2 exhibiting the greatest accumulation (Figure 4D). A similar pattern was observed for N RNA, where APOE2/2 and APOE4/4 showed elevated levels compared to APOE3/3, though the difference between APOE2/2 and APOE4/4 did not reach statistical significance (Figure 4E). These data suggest that APOE2 and APOE4 may enhance viral replication or impair viral clearance mechanisms.

To explore genotype-specific molecular consequences, we implemented an integrative analytic workflow to identify SCV2-responsive genes modulated by APOE isoforms (Figure 4F). Differential expression analysis revealed distinct transcriptional landscapes. In APOE2/3 versus APOE3/3 comparisons, we observed significant alterations in innate immune, interferon-responsive, and vascular signaling genes (Figure 4G). Comparisons between APOE2/3 and APOE4/4 revealed divergent regulatory patterns (Figure 4H–I), indicating that APOE isoforms may modulate host susceptibility through distinct molecular pathways.

We next evaluated whether APOE genotype affected the expression of known viral entry and antiviral response factors. Cathepsin L (CTSL), a lysosomal protease involved in spike protein priming, was modestly upregulated in APOE4/4 at both the mRNA (Figure 4J) and protein levels (Figure 4K) under baseline (uninfected) conditions. APOE2/2 also showed elevated CTSL protein, albeit to a lesser extent than APOE4/4. FURIN, another spike-processing protease, was significantly elevated in APOE4/4 relative to APOE3/3, with a trend toward further induction post-infection (Figure 4L). Notably, ACE2, the primary SCV2 receptor, was significantly upregulated in APOE2/2 compared to APOE3/3 (Figure 4M), suggesting both APOE2 and APOE4 isoforms may increase susceptibility through enhanced entry factor expression.

Expression of interferon receptors IFNAR1 and IFNAR2 was markedly reduced in APOE3/3 compared to both APOE2/2 and APOE4/4 (Figure 4N–O), indicating genotype-dependent disparities in baseline antiviral signaling capacity that could influence downstream immune responses.

To validate and expand upon these findings, we profiled additional SCV2 entry and immune-regulatory genes across APOE genotypes (Figure S4B–F). While genes such as TMPRSS2, NLRP1, TPCN3, and RAB14 showed no significant genotype-specific differences, TMPRSS4 trended higher in APOE2/2 (p = 0.113), and CTSL was significantly elevated in APOE4/4 relative to both APOE3/3 and APOE2/2—supporting a gradient of entry factor expression favoring APOE4.

A schematic overview of the SCV2 viral lifecycle contextualized these observations (Figure S4G), and a summary table (Figure S4H) consolidated the genotype-dependent trends in host susceptibility factors. Both APOE2/2 and APOE4/4 exhibited elevated expression of ACE2, CTSL, FURIN, and IFNAR1/2, while APOE3/3 consistently demonstrated lower baseline expression, consistent with a comparatively “resilient” transcriptional profile.

Together, these findings reveal a bidirectional and context-specific influence of APOE isoforms on viral RNA burden, host factor expression, and interferon signaling. The parallel susceptibility patterns observed in APOE2/2 and APOE4/4 FORMA-COs suggest that, despite their contrasting roles in Alzheimer’s disease risk, these isoforms may converge on shared molecular pathways that exacerbate neuroimmune dysfunction in the setting of SCV2 infection.

### APOE Genotype-Specific Regulation of VEGF Signaling and Angiogenic Pathways Following SCV2 Infection

To delineate how APOE genotype modulates immune-vascular responses during SCV2 infection, we conducted KEGG pathway enrichment analysis of DEGs derived from two publicly available datasets: bulk RNA-seq of postmortem human brains (GSE144254) and single-cell RNA-seq from postmortem brain tissue of ROSMAP participants. This analysis revealed genotype-specific alterations in angiogenic, metabolic, and immune-related signaling pathways, with particular emphasis on VEGF signaling—a critical regulator of endothelial function, vascular remodeling, and inflammation.

Comparisons between APOE2/3 and APOE3/3 revealed widespread transcriptional reprogramming across multiple biological systems (Figure 5A). Notably enriched pathways included VEGF signaling, T cell receptor signaling, PI3K-Akt signaling, and PD-L1/PD-1 checkpoint pathways, implicating APOE2/3 in modulating immune activation and vascular remodeling. Additional pathways involved cell cycle regulation, p53 signaling, osteoclast differentiation, and neurodegenerative mechanisms such as ALS and axon guidance, suggesting broad systemic influence of APOE2/3 beyond antiviral immunity.

**Figure 5.**
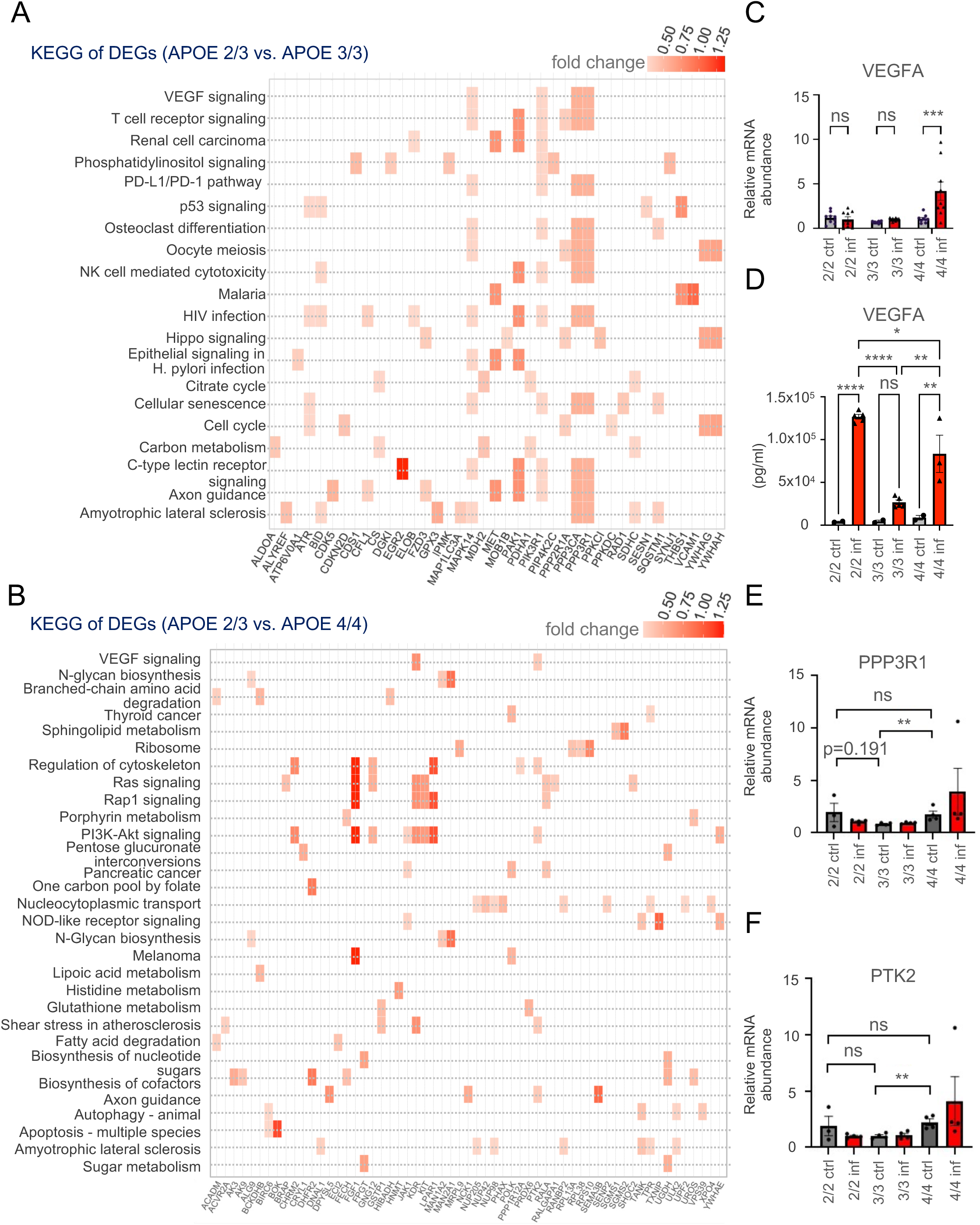
**APOE Genotype Modulates Angiogenic, Immune, and Metabolic Pathways via VEGF Signaling.** (A–B) Heatmaps of KEGG pathway enrichment for DEGs comparing APOE2/3 vs. APOE3/3 (A) and APOE2/3 vs. APOE4/4 (B). Notable pathways include VEGF, PI3K-Akt, and PD-1/PD-L1 signaling, along with metabolic, apoptotic, and neurodegenerative pathways. (A) Bar graph showing VEGFA mRNA expression. Significant induction observed in APOE4/4-infected cells (***p < 0.001); no changes in APOE2/2 or APOE3/3. (B) VEGFA protein levels show highest upregulation in APOE2/2, followed by APOE4/4; minimal change in APOE3/3. (C) PPP3R1 mRNA expression is significantly elevated in APOE4/4 at baseline (***p < 0.001), with no changes post-infection. (D) PTK2 mRNA levels are significantly higher in APOE4/4 controls (**p < 0.01), unchanged in 2/2 or 3/3 groups.

In contrast, comparison between APOE2/3 and APOE4/4 highlighted distinct transcriptional signatures (Figure 5B). APOE4/4 organoids exhibited upregulation of angiogenesis-associated pathways, including VEGF, Ras, Rap1, and PI3K-Akt signaling, alongside elevated expression of genes linked to sphingolipid metabolism, N-glycan biosynthesis, autophagy, and apoptosis.

These results suggest that APOE4/4 is transcriptionally primed for stress-adaptive, angiogenic, and metabolic reprogramming, while APOE2/3 maintains a comparatively homeostatic state.

Focusing on the VEGFA axis, we observed robust genotype-specific regulation. Although VEGFA mRNA expression remained low in APOE2/2 and APOE3/3 organoids, infection triggered a significant increase in APOE4/4 (Figure 5C, ***p < 0.001), indicating an infection-inducible angiogenic response unique to the APOE4 genotype. Surprisingly, VEGFA protein levels—quantified by multiplex MSD assay—were highest in APOE2/2 organoids following infection, followed by APOE4/4, with minimal induction in APOE3/3 (Figure 5D). This disparity between transcript and protein levels suggests post-transcriptional regulation of VEGFA in APOE2/2, revealing a previously unrecognized mechanism of vascular modulation.

Expression of PPP3R1, a calcineurin subunit involved in calcium signaling and endothelial remodeling, was significantly elevated at baseline in APOE4/4 compared to APOE3/3 (Figure 5E, ***p < 0.001). Likewise, PTK2 (focal adhesion kinase), a key mediator of VEGF-induced endothelial activation, was upregulated in APOE4/4 under control conditions (Figure 5F, **p < 0.01), suggesting a vascular-primed transcriptional state associated with APOE4.

To further dissect the mechanisms by which APOE genotype influences immune-vascular remodeling during SCV2 infection, we mapped KDR, MAPK14, PIK3R1, and PPP3CA. Although KDR (VEGFR2)—the primary receptor for VEGFA—did not differ significantly across genotypes, we observed a trend toward increased expression in APOE2/2 controls (p = 0.079), along with a slight post-infection increase in APOE4/4 organoids (Figure S5B). This suggests potential enhancement of VEGF receptor signaling in both genotypes under distinct conditions.

Downstream components also showed modest genotype-linked trends. MAPK14 (encoding p38 MAPK) exhibited a nonsignificant increase in APOE4/4-infected FORMA-COs (Figure S5C), while PIK3R1, a regulatory subunit of PI3K, trended upward in APOE2/2 controls and APOE4/4-infected groups (p = 0.184; Figure S5D). Additionally, PPP3CA—the catalytic subunit of calcineurin—was slightly elevated in APOE4/4 post-infection, though this change did not reach statistical significance (Figure S5E).

Collectively, these patterns suggest that APOE4/4 FORMA-COs exhibit coordinated upregulation of pro-angiogenic signaling through VEGFA transcription and select downstream effectors. In contrast, APOE2/2 appears to drive VEGF protein accumulation via potential post-transcriptional mechanisms, with only modest changes at the mRNA level, suggesting mechanistically distinct but functionally convergent vascular remodeling in response to SCV2 infection across different APOE genotypes.

### SCV2 Infection Drives Genotype-Dependent Inflammatory, Vascular, and Glial Alterations in APOE iPSC-Derived and hACE2 Mouse Brain Models APOE Isoforms Differentially Regulate Pro- and Anti-inflammatory Signaling Pathways

To examine how APOE genotype modulates cytokine responses and neurovascular remodeling during viral insult, we quantified inflammatory protein levels in isogenic APOE2/2, APOE3/3, and APOE4/4 FORMA-COs using multiplexed MSD assays and performed in vivo validation in hACE2 transgenic mice infected with SCV2.

IL-1α, a proinflammatory cytokine linked to neuroinflammation, was significantly elevated in APOE4/4 organoids following infection, while APOE2/2 and APOE3/3 organoids exhibited only modest, nonsignificant changes (Figure 6A). Interestingly, baseline IL-1α levels were higher in APOE2/2 and APOE3/3, suggesting that APOE4 may potentiate infection-specific IL-1α upregulation. Conversely, TNF-β was robustly induced in APOE2/2 organoids upon infection, exceeding levels seen in APOE3/3 and APOE4/4 (Figure 6B). This divergence points to genotype-specific inflammatory programs: APOE4 may amplify IL-1α–mediated responses, while APOE2 preferentially activates TNF-β signaling under stress.

**Figure 6.**
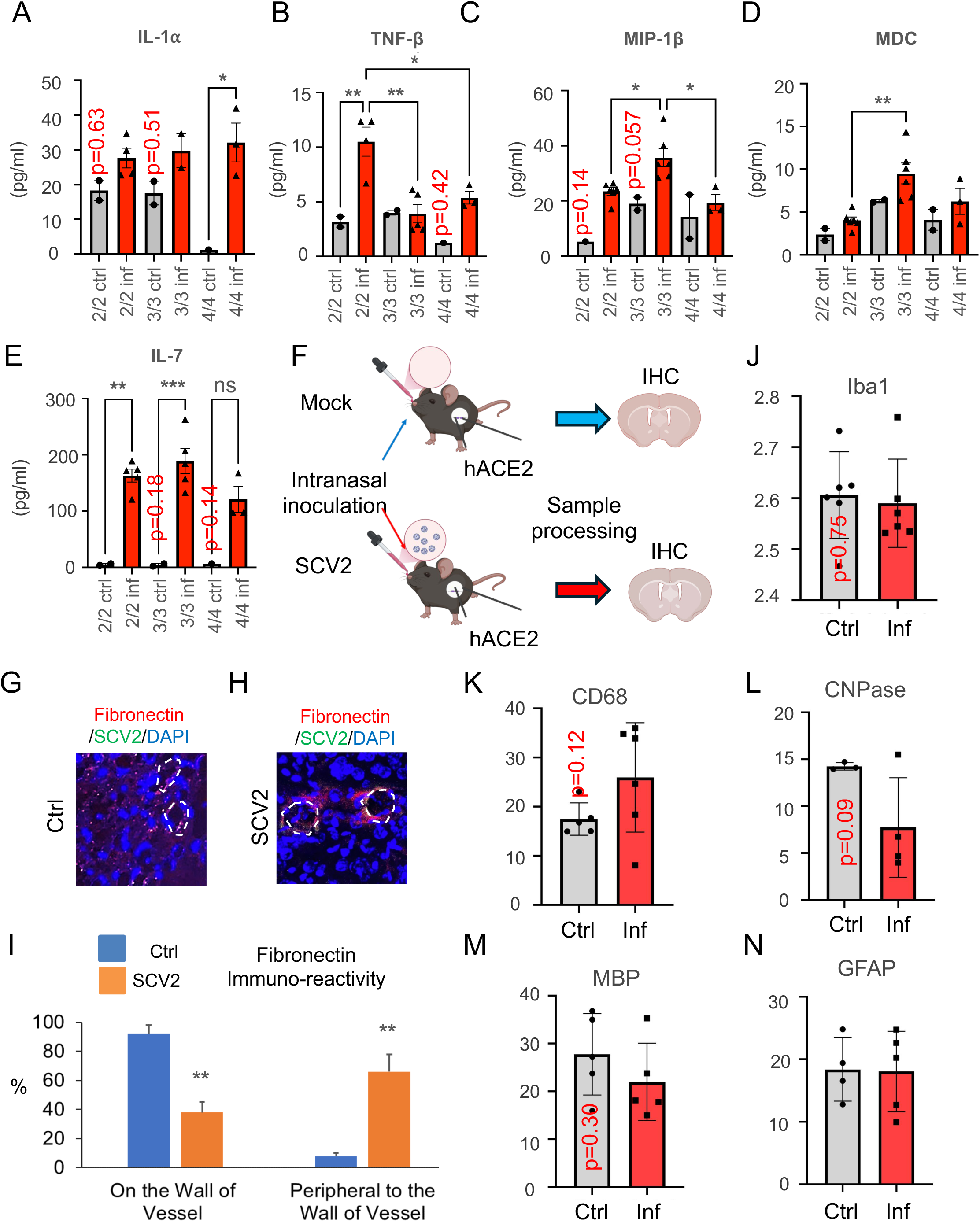
**APOE Genotype Modulates Inflammatory, Vascular, and Glial Responses to SARS-CoV-2.** (A–E) MSD quantification of cytokines across APOE genotypes (2/2, 3/3, 4/4) under mock (gray) and infection (red) conditions. APOE4/4 cells show increased IL-1α; APOE2/2 cells show elevated TNF-β; APOE3/3 cells trend toward higher MIP-1β and MDC; IL-7 is reduced in APOE4/4 upon infection. (A) Schematic of hACE2 mouse model with nasal SARS-CoV-2 application. (G–I) Immunofluorescence showing increased perivascular fibronectin in infected brains, suggesting extracellular matrix remodeling. Quantification shows decreased vessel wall signal and increased perivascular signal post-infection. (J–K) CD68+ microglial activation increases in infected mouse brains (non-significant); Iba1 expression unchanged. (L–M) CNPase expression trends lower with infection; MBP shows no change. (N) GFAP levels unchanged across conditions.

MIP-1β levels rose across all genotypes post-infection, with the strongest induction in APOE3/3 and weakest in APOE4/4 (Figure 6C). MDC, a chemokine involved in leukocyte recruitment, was significantly elevated in APOE3/3 organoids but remained unchanged in APOE2/2 and APOE4/4 (Figure 6D). IL-7, essential for lymphocyte maintenance and T cell survival, was markedly suppressed in APOE4/4 after infection (Figure 6E), suggesting a genotype-linked impairment of adaptive immune support.

To further dissect genotype-specific immune modulation, we expanded cytokine profiling across additional inflammatory mediators (Figure S6). IL-8, a neutrophil chemoattractant, was most elevated in APOE3/3 post-infection, while APOE4/4 showed blunted induction (Figure S6A). IL-12, a Th1-promoting cytokine, was modestly upregulated in APOE3/3 and APOE4/4 (Figure S6B), whereas IL-15, critical for NK and CD8⁺ T cell responses, was highest in APOE4/4 (Figure S6C). In contrast, IL-16, a chemoattractant and T cell modulator, showed greater induction in APOE2/2 (Figure S6D). Other chemokines displayed similar patterns: IP-10 was preferentially elevated in APOE4/4; MCP-4 increased broadly but was highest in APOE2/2 and APOE4/4 (Figure S6E–F); and Eotaxin was strongly induced in APOE3/3, followed by APOE4/4 (Figure S6G). MIP-1α reached peak levels in APOE4/4-infected organoids (Figure S6H).

A summary of cytokine expression (Figure S6I) revealed that APOE4/4 is biased toward pro-inflammatory and chemotactic signaling, APOE2/2 emphasizes TNF and IL-16 responses, and APOE3/3 maintains a relatively balanced cytokine profile. These patterns suggest evolutionarily distinct immunological strategies encoded by each APOE isoform—APOE4 enhancing acute inflammation, APOE2 favoring TNF-centered immunity, and APOE3 supporting immune equilibrium.

To validate these findings in vivo, we used a nasal instillation model of SCV2 infection in hACE2 transgenic mice (Figure 6F). Immunohistochemical analysis revealed fibronectin redistribution from vascular walls to perivascular spaces in infected brains (Figure 6G–I), indicating vascular barrier disruption and extracellular matrix remodeling. Elevated CD68 expression—an indicator of microglial activation—was observed in infected brains compared to mock controls (Figure 6J– K), although not statistically significant. CNPase, a marker of oligodendrocyte function, was modestly reduced following infection (p = 0.09), whereas MBP and GFAP levels remained unchanged (Figure 6L–N), suggesting early or subclinical glial perturbations that may precede overt demyelination or astrogliosis.

Collectively, these results demonstrate that APOE genotype modulates inflammatory responses, vascular remodeling, and glial vulnerability during SCV2 infection, both in human iPSC-derived organoid models and in vivo mouse systems.

## Discussion

Our study demonstrates that common human APOE variants confer distinct susceptibilities to neuroimmune and vascular injury under pathogenic stress. Using isogenic human iPSC-derived immune-vascularized forebrain organoids (FORMA-COs), coupled with in vivo hACE2 mouse validation, we show that APOE2 and APOE4 alleles orchestrate divergent yet convergent responses to SCV2 infection. While APOE3/3 appears relatively resilient, APOE2/2 and APOE4/4 exhibit exaggerated inflammatory and vascular perturbations, highlighting APOE genotype as a context-dependent regulator of host responses to environmental insults.

FORMA-COs revealed that both APOE2/2 and APOE4/4 organoids accumulate significantly more viral RNA than APOE3/3, suggesting isoform-dependent deficits in viral clearance or differential permissiveness to infection. However, their downstream molecular responses diverged. APOE4/4 organoids mounted a strong IL-1α–driven inflammatory program and activated angiogenic mediators including VEGFA, PTK2, and PPP3R1, suggesting a maladaptive endothelial remodeling response. In contrast, APOE2/2 organoids showed increased TNF-β production and elevated VEGFA protein levels despite limited transcript induction, implying post-transcriptional or metabolic regulation. These distinct cytokine and vascular signatures were mirrored in SCV2-infected hACE2 mouse brains, which exhibited APOE-linked fibronectin redistribution, microglial activation, and early glial stress.

These findings expand the known roles of APOE beyond lipid metabolism and Alzheimer’s disease risk. While APOE4 is widely recognized as a major genetic risk factor for late-onset AD and cerebrovascular pathology, our data suggest that APOE4 also predisposes the neurovascular unit to exaggerated inflammatory and angiogenic responses following acute viral exposure. This aligns with prior studies linking APOE4 to blood-brain barrier disruption, endothelial dysfunction, and increased vulnerability to hypoxia or infection^1–7^.

Importantly, our study also highlights APOE2 as a genotype conferring non-trivial risk. Although historically considered protective against AD, APOE2 has been associated with cerebrovascular anomalies, dysregulated lipid trafficking, and type III hyperlipoproteinemia^8–14^. In our system, APOE2/2 organoids exhibited unique transcriptional and cytokine profiles, including strong TNF-β signaling, altered VEGF regulation, and elevated accumulation of viral RNA. These findings suggest that APOE2 may sensitize the brain to inflammatory and vascular stress via distinct, isoform-specific mechanisms—challenging the simplistic protective-versus-pathogenic dichotomy of APOE isoforms.

The FORMA-CO platform provided a powerful model to dissect these isoform effects across multiple brain-relevant cell types. The inclusion of neurons, astrocytes, oligodendrocyte-lineage cells, endothelial cells, pericytes, and microglia within an isogenic background enabled a high-resolution view of genotype-specific host-pathogen interactions. Notably, APOE4/4 organoids showed diminished IL-7 production and reduced expression of lymphoid-supportive cytokines after infection, suggesting impaired adaptive immune support consistent with immune senescence^52–54^ and poor outcomes in older APOE4 carriers^55–61^.

In vivo experiments in hACE2 mice further supported these findings, revealing early APOE-dependent changes in vascular matrix integrity and glial activation—even in the absence of overt neurodegeneration. These subclinical alterations parallel those observed in aging and early neurodegenerative states, suggesting that APOE may prime the brain for long-term injury in response to repeated or unresolved stress^55–62^.

Collectively, our data establish that APOE genotype shapes the trajectory of neurovascular and immune responses to SCV2 infection through distinct isoform-specific programs. These include differences in viral burden, cytokine signaling, angiogenic remodeling, and immune regulation. While SCV2 served as a prototypical pathogenic stressor, the implications of our findings likely extend to other infectious, metabolic, and age-related insults. This work thus provides a mechanistic framework for understanding how APOE genotype influences individual vulnerability to neuroinflammation and vascular dysfunction—key processes implicated in long-COVID, vascular dementia, and neurodegenerative disease progression^63^.

### Study limitations and implications

While our integrated organoid and in vivo systems provide valuable insights, several limitations should be noted. First, FORMA-COs, though complex, do not fully replicate systemic physiology—particularly lacking peripheral immune components, long-range vascular perfusion, and hormonal signaling. As such, chronic, multisystem interactions remain outside the scope of this model. Second, although our iPSC lines are isogenic with respect to APOE, they derive from a single donor background; thus, additional donor-matched or population-diverse lines will be needed to confirm generalizability. Third, we focused on early-phase responses to a single stressor— SCV2 infection—leaving longer-term consequences and cross-stressor comparisons to future work.

Future studies should explore how APOE genotype interacts with other genetic modifiers (e.g., *TREM2*, *ACE2*, *IFNAR1*), metabolic status, or repeated immune challenges. Incorporating vascular perfusion systems, longitudinal stress paradigms, and spatial omics technologies may further illuminate how APOE shapes adaptive versus maladaptive trajectories in the human brain. Importantly, our findings offer translational implications: individuals with APOE2 or APOE4 genotypes may benefit from therapies targeting VEGF, IL-1, or IFN pathways to prevent or reverse neurovascular injury under inflammatory stress.

These insights are likely to extend beyond SCV2 to broader conditions such as post-viral cognitive impairment, long COVID, and sporadic neurodegenerative diseases. By establishing a mechanistic framework for APOE-mediated injury in a human neurovascular context, our study lays the groundwork for genotype-informed therapeutic strategies that address both immediate and long-term brain health risks.

## Materials and Methods

### Ethics statement

All work involving infectious SARS-CoV-2 was conducted under biosafety level 3 (BSL-3) and animal BSL-3 conditions, as approved by the Johns Hopkins Institutional Biosafety Committee (IBC Protocol #P2003270301). SARS-CoV-2 inactivation protocols were validated and approved for specimen removal from high-containment laboratories. All animal procedures conformed to the National Institutes of Health Guide for the Care and Use of Laboratory Animals and were approved by the Johns Hopkins University Institutional Animal Care and Use Committee (IACUC Protocols: MO19M422, MO19M98, MO18M58).

### Animal Experiments

Heterozygous K18-hACE2 C57BL/6J mice (strain: 2B6.Cg-Tg(K18-ACE2)2Prlmn/J; The Jackson Laboratory) were bred and maintained at Johns Hopkins University School of Medicine. Mice (6–8 weeks old; n=8 per genotype, balanced by sex) were housed in groups of 1–5 and provided standard chow. Mice were intranasally inoculated with 25 μL of Dulbecco’s Modified Eagle Medium (DMEM) containing 2.5 × 10⁴ PFU of SARS-CoV-2 (USA-WA1/2020 strain) or vehicle (DMEM alone; n=3 per sex, mock controls). Body weights were monitored daily. At 6 days post-infection, mice were euthanized via isoflurane overdose, and tissues were harvested following cardiac perfusion with 2 mL PBS. Lungs were collected for viral RNA quantification, and brains/olfactory bulbs were fixed for immunohistochemical analysis.

### Human iPSC Culture and APOE Isogenic Line Derivation

Human induced pluripotent stem cell (iPSC) lines homozygous for APOE2/2, APOE3/3, or APOE4/4 were generated using CRISPR/Cas9 editing at the rs429358 and rs7412 loci. The parental APOE4/4 BC1 iPSC line was originally derived by Dr. Linzhao Cheng (Johns Hopkins University) from CD34⁺ human bone marrow cells via episomal reprogramming with a non-integrating plasmid system. iPSCs were cultured in DMEM/F12 (Gibco, Cat# 11330-032) supplemented with 20% KnockOut Serum Replacement (Gibco, Cat# 10828-028), 4 ng/mL FGF2 (PeproTech, Cat# 100-18B), 1 mM GlutaMAX (Gibco, Cat# 35050-061), 100 μM non-essential amino acids (Gibco, Cat# 11140-050), and 100 μM 2-mercaptoethanol (Sigma-Aldrich, Cat# M3148). Medium was changed daily, and cells were passaged every 4–6 days using 1 mg/mL collagenase type I (Gibco, Cat# 17100-017) in DMEM/F12 at a 1:6 to 1:12 split ratio. All procedures involving human pluripotent stem cells were conducted under the oversight of the Johns Hopkins Institutional Stem Cell Research Oversight (ISCRO) Committee.

### Derivation, Culture, and Tri-linage Differentiation of Human FOXG1⁺ Forebrain Neural Progenitor Cells

To initiate neural differentiation, human ESC or iPSC colonies were pre-treated with 1 mg/mL collagenase type I (Gibco, Cat# 17100-017) in DMEM/F12 for 5–10 minutes at 37°C. As colony borders began to lift while centers remained attached, collagenase was removed and replaced with growth medium. Detached colonies, free of underlying MEFs, were carefully transferred to low-attachment six-well plates (Corning, Cat# 3471) and cultured in suspension for 2 days in human ESC medium lacking FGF2. From days 2 to 6, dual-SMAD inhibition was initiated by supplementing the medium (hereafter referred to as KoSR medium) with 50 ng/mL recombinant human Noggin (R&D Systems, Cat# 6057-NG-025), 1 μM dorsomorphin (Tocris, Cat# 3093), and 10 μM SB431542 (Tocris, Cat# 1614). On day 7, the resulting embryoid bodies (EBs) were transferred to plates coated with either Matrigel (Corning, Cat# 354277) or laminin (Sigma-Aldrich, Cat# L2020) and cultured in N2 induction medium composed of DMEM/F12 (Gibco, Cat# 11330-032), 1% N2 supplement (Gibco, Cat# 17502-048), 100 μM MEM non-essential amino acids (Gibco, Cat# 11140-050), 1 mM GlutaMAX (Gibco, Cat# 35050-061), and 2 μg/mL heparin (Sigma-Aldrich, Cat# H3149). Cultures were fed with N2 medium every other day from days 7 to 12, and daily thereafter. By days 8–9, attached EBs flattened and differentiated into neural rosettes. Continued induction led to the emergence of compact, columnar 3D neural aggregates—termed rosette-derived neural aggregates (RONAs)—in the centers of rosette colonies. These RONAs were manually microdissected with care to avoid contaminating peripheral flat cells or underlying layers, and maintained as neurospheres in Neurobasal medium (Gibco, Cat# 21103-049) supplemented with B27 minus vitamin A (Gibco, Cat# 12587-010) and 1 mM GlutaMAX for 24 hours. RONAs were then enzymatically dissociated into single cells and replated onto laminin/poly-D-lysine–coated plates (Sigma-Aldrich, Cat# P0899) for downstream assays.

Astrocyte differentiation (Adapted from Perriot et al., 2021)^64^, Day-10 neural rosettes were transferred to Glial Expansion Medium to derive glial progenitor cells (GPCs), which were expanded for ≥8 passages and cryopreserved. For astrocyte differentiation, GPCs were thawed and plated on Matrigel-coated flasks at 50,000 cells/cm² in Astrocyte Induction Medium for 14 days, then switched to Astrocyte Maturation Medium containing CNTF (20 ng/mL, PeproTech) for 4 weeks. Cultures were passaged as needed to prevent overgrowth. Astrocytes began to emerge by day 5–7 and progressively enriched over time. CNTF was withdrawn after maturation, and cells were maintained in Astrocyte Maintenance Medium. Differentiation was repeated 2–3 times per iPSC line to ensure reproducibility.

Oligodendrocyte differentiation (Adapted from Wang et al., 2013)^65^. Human iPSCs or ESCs were dissociated and aggregated into embryoid bodies (EBs), which were sequentially patterned in neural induction medium with bFGF, heparin, and retinoic acid. From day 14–28, colonies were exposed to purmorphamine and B27 to promote neural epithelial and pre-OPC lineage specification. Cells were assessed for OLIG2 and NKX2.2 expression, then transitioned to glial induction medium containing T3, biotin, cAMP, PDGF-AA, IGF-1, and NT3. Resulting gliospheres were cultured for up to 200 days, yielding oligodendrocyte progenitor cells (OPCs).

### Differentiation and Characterization of iPSC-Derived Human Yolk Sac–Like Tissues

Human iPSCs were differentiated into yolk sac–like tissue using a modified protocol that mimics early extra-embryonic hematopoiesis and vasculogenesis pathways, giving rise to both floating macrophage-like cells (iMacs) and adherent endothelial (iEndo) and pericytic (iPeri) cells localized along the basement membrane. We differentiate iPSC to microglia from yolk sac precursors by inducing a process reminiscent of yolk sac progenitors^66, 67^. Basal microglia differentiation medium (MDM) consists of Stempro-34 SFM supplemented with transferrin Ferric ammonium citrate (FAC) (50 μM), glutamic acid (2 mM), ascorbic acid (0.5 mM) and MTG (0.45 mM). 75% confluent iPS were passaged on to a Matrigel-coated 6-well plate at a density of 1.0 × 10^5^ cells/well. Starting from the next day after plating (defined as day 0, d0), full media were changed every other day, the cells were cultured for the next 16 days in MGM supplemented with the following cytokines: *<u>d0</u>*, BMP4 (5 ng/mL), VEGF (50 ng/mL), and TWS119 (2 μM); *<u>d2</u>*,

BMP4, VEGF, and FGF2 (20 ng/mL); *<u>d4</u>*, VEGF and FGF2; *<u>d6-10</u>*, VEGF, FGF2 (50 ng/mL), ISCK03 (2 μM), IWR-1 (5 μM), IL-6 (10 ng/mL), and IL-3 (20 ng/mL); *<u>d12-14</u>* FGF2, SCF, IL-6, and IL-3; *<u>d16-25</u>*, CSF-1 (50 ng/mL). Floating cells typically appeared around day 7 and they were re-plated back to the same culture vessels during medium changes. Twenty-five days post differentiation, floating microglia progenitors were maintained in modified microglia growth medium (MGM)^68^consists of human neuronal conditioned DMEM:F12 (1:1) medium supplied with glutamic acid, N-acetyl cysteine (5 µg/mL), insulin (5 µg/mL), FAC (50 μM), sodium selenite (100 ng/mL), TGF-β2 (2 ng/mL), CXCL12 (100 ng/mL), IL-34 (100 ng/mL), cholesterol (1.5 µg/mL) and oleic acid (0.1 µg/mL), gondoic acid (0.001 µg/mL) and heparan sulfate (1 µg/mL, Galen Laboratory Supplies) on collagen IV (2 µg/mL) coated plates at a density of 3×10^4^/cm^2^. All chemicals or growth factors were from Sigma or Peprotech unless otherwise noted. Two weeks after MGM maintenance, microglia cells were used for experiments.

### Human iMac, iEndo, and iPeri Migrate and Integrate into Pre-patterned Cerebral Organoids

Immune-Vascularized forebraind organoid generation was performed in three sequential steps: (1) induction of forebrain-patterned organoids from human iPSC-derived FOXG1⁺ NPCs; (2) parallel differentiation of yolk sac–like progenitors into macrophage-like (iMac), endothelial (iEndo), and pericytic (iPeri) cell populations; and (3) timed co-embedding of these supporting cells into the organoid matrix.

Different human iPS lines have variable neural differentiation potency, which often lead to compromised robustness of organoid system^69^. To increase cellular and structural consistency, we use the defined and enriched multipotent FOXG1-hNPC for generation of cerebral organoids. This strategy is different from most of the conventional methods that use embryonic bodies as starting materials for creating brain organoids, we anticipate the improvement will increase the organoid uniformity that are crucial for the functional comparison. FOXG1-NSC were dissociated into single cells and loaded into each well of the low-attachment 96-well plates (round bottomed, Corning) with defined number (30,000, 45,000 and 60,000), a forced quick aggregation centrifugation^70^ of low-attachment 96-well plate (200g, 5 mins) was applied. One week post the forced aggregation, each FOXG1 neurosphere was embedded within a single droplet Matrigel (7 μl) one well of Organoid Embedding Sheet (STEMCELL Technologies).

Three weeks after the formation of FOXG1-prepatterned organoids (analogous to the brain rudiment), the organoids were transferred to the well of Organoid Embedding Sheet and embedded in a single droplet (20 μl) containing 30,000 iPS derived human iMacs, iEndo, and iPeri. The organoids were positioned in the middle of Matrigel droplet before it gets solidified. The droplets were cultured in differentiation medium for 7 more days and then transferred to an orbital shaker with medium change every 5-7 days. Neural differentiation medium contains astrocyte-conditioned neurobasal/B27, BDNF (20ng/ml), GDNF (20 ng/ml), ascorbic acid (0.2 mM), dibutyryl cAMP (0.5 mM), N-acetyl cysteine (5 µg/mL), insulin (5 µg/mL), FAC (50 μM), sodium selenite (100 ng/mL), TGF-β2 (2 ng/mL), CXCL12 (100 ng/mL), IL-34 (100 ng/mL), cholesterol (1.5 µg/mL) and oleic acid (0.1 µg/mL), gondoic acid (0.001 µg/mL) and heparan sulfate (1 µg/mL, Galen Laboratory Supplies). To generate a balanced excitatory and inhibitory neuron subtypes, 7 days post-embedding, the organoids was administrated with retinoic acid (2 µM) for 7 days^37^. The organoids were sectioned (10µm). Diverse cell types were analyzed.

### Measure the sphericity of neurospheres

The measurement of FOXG1 neurosphere sphericity was carried out using Zeiss Axiovert Inverted Phase Contrast Fluorescence Microscope combined with image analysis software. To begin, FOXG1 neurospheres are cultured under appropriate conditions, in low-adhesion plates, and then harvested for imaging. High-resolution images of individual neurospheres are captured using an inverted microscope, ensuring the entire neurosphere is in focus. These images are then imported into image analysis software ImageJ. In the software, the images are processed to enhance contrast, often by converting them to grayscale and applying thresholding to distinguish the neurosphere from the background. Edge detection methods are then used to outline the perimeter of each neurosphere. The major axis (longest diameter) and minor axis (shortest diameter) are measured, and the sphericity is calculated using the formula: Sphericity= 2⋅ (radius of the sphere)/ (major axis length + minor axis length). For a perfectly spherical neurosphere, the sphericity value should approach 1, with lower values indicating deviations from a spherical shape. Multiple neurospheres were analyzed to obtain reliable average sphericity values, and statistical comparisons can be made across different experimental conditions if needed. Consistency in imaging parameters and software calibration is crucial to ensure accurate measurements.

## Cells and Viruses

Vero C1008 cells [Vero 76, clone E6, Vero E6] (ATCC CRL-1586) were obtained from the American Type Culture Collection (ATCC, Manassas, VA, USA) and were used to generate virus and determine infectious virus titers. Vero C1008 cells were maintained in Dulbecco’s Modified Eagle Medium (DMEM) supplemented with 10% FBS, 1mM L-glutamine, 50U/mL penicillin and 50 ug/mL streptomycin. SARS-CoV-2/Wuhan-1/2020 virus (U.S. Centers for Disease Control and Prevention) was provided by Dr. Andrew Pekosz. 2019-nCoV/USA-WA1/2020 was obtained through BEI Resources, National Institute of Allergy and Infectious Diseases (NIAID), National Institutes of Health (NIH).

### Generation and Quantification of SCV2 Virus Stocks

Generation of P1 and working stocks of SARS-CoV-2 was performed in Vero C1008 cells inoculated with virus at a multiplicity of infection 0.01 in DMEM supplemented with 2% fetal bovine serum, 1 mM L-glutamine for 2 d. Supernatants were collected, centrifuged at 500 x g for 5 minutes, aliquoted, and stored at −80degC. Infectious virus titers were determined by tissue culture infectious dose 50 (TCID_50_) assay using Vero C1008 cells as previously described (Bullen CK, Davis SL, Looney MM. Quantification of Infectious SARS-CoV-2 by the 50% tissue culture infectious dose endpoint dilution assay. Methods Mol Biol. In press).

### SCV2 infection of Brain Assembloids

Brain assembloids were infected with SARS-CoV-2/Wuhan-1/2020 virus (MOI 0.1) in a 48-well plate containing 10 assembloids per well and rocked gently every ten minutes for the first hour of inoculation to ensure homogenous virus distribution. Six hours post infection, brain assembloids were washed two times with one half-volume changes of PBS and provided a 1:1 ratio mix of fresh and culture conditioned neuronal growth media. One half of the culture media was replaced with fresh neuronal growth media every three days until the final time point at nine days post infection. At indicated time points, supernatant samples were collected for viral quantification and cytokine analysis, and assembloids were either formaldehyde fixed for immunohistochemistry or lysed for RNA extraction, viral RNA quantification and gene expression analysis.

### RNA Extraction and Viral RNA Quantification

Zymo Quick-RNA Viral 96 Kit was used to isolate RNA from supernatants according to the manufacturer’s protocol (Zymo Research). For total RNA extraction from brain assmbloids or mouse lung tissue, samples were homogenized in 1 mL TRIzol reagent with ceramic 1.4 mm beads for 2 cycles of 30 seconds at 6800 rpm (Precelley’s Evolution). Total RNA from mouse lung tissue was extracted using chloroform phase separation and further purified using the RNeasy Mini Kit with on-column DNase digestion per the manufactures protocol (Qiagen). Total RNA from brain assembloids was extracted using chloroform phase separation and isopropyl alcohol precipitation and). RNA was subsequently DNase digested using TURBO DNase and reextracted using acid-phenol:choloroform, pH 4.5 (Ambion) per the manufacturer’s protocol (Ambion).

cDNA was synthesized using qScript cDNA Supermix containing random hexamers and oligo-dT primers (Quanta Biosciences). Real-time PCR was performed in technical triplicate for each sample using TaqMan Fast Advanced Master Mix (Applied Biosystems) on an on a StepOne Plus Real Time PCR machine (Applied Biosystems). SARS-CoV-2 RNA was detected using the following primers and probe targeting the region of the SARS-CoV-2 nucleocapsid (N) gene (COVID-19 CDC Research Use Only kit, Integrated DNA Technologies, Catalog #10006713): forward (5’-TTACAAACATTGGCCGCAAA-3’), reverse (5’-GCGCGACATTCCGAAGAA-3’) and probe (5’-FAM-ACAATTTGCCCCCAGCGCTTCAG-BHQ1-3’). The cycling parameters were as follows: (i) 2 min at 50 °C; (ii) 2 min at 95 °C; and (iii) 45 cycles at 95 °C for 3 s and 55 °C for 30 s. Serial dilutions of a plasmid containing the SARS-CoV-2 N gene were used to generate molecular standard curves (Integrated DNA Technologies, Catalog #10006625). Viral RNA load in mouse lung tissue was adjusted for tissue weight. Viral RNA copies per μg of total RNA from brain assembloids were normalized to the human RNA Polymerase II gene (Pol2Ra) using the TaqMan gene expression assay (Hs00172187_m1; Applied Biosystems).

### Infectious Virus Quantification

Supernatants from brain assembloid cultures were collected for virus titer by the TCID_50_ assay as described previously (Bullen CK, Davis SL, Looney MM. Quantification of Infectious SARS-CoV-2 by the 50% tissue culture infectious dose endpoint dilution assay. Methods Mol Biol. In press). In brief, supernatants from the brain assembloids were serially diluted in seven-point half-log dilutions and incubated on Vero C1008 cells in six technical replicates, leaving the bottom row uninfected as a control. Each supernatant sample was measured in triplicate. Plates were scored for CPE five days post infection using the CellTiter-Glo® Luminescent Cell Viability Assay per the manufacture’s protocol (Promega). TCID_50_ was calculated using the Reed and Muench method (Lindenbach BD. Measuring HCV infectivity produced in cell culture and in vivo. *Methods Mol Biol*. 2009;510:329-336. doi:10.1007/978-1-59745-394-3_24).

### Cytokine Analysis

Supernatants from brain assembloid cultures were collected at indicated time points and cytokines were measured using the V-PLEX neuroinflammation panel 1 (K15210D) and U-PLEX interferon combo (K15094K) from Meso Scale Diagnostics according to the manufacture’s protocol. Data were acquired on a Meso Quickplex SQ120 and analyzed using …

### Nanostring Gene Expression Analysis

Total RNA purified from mock-infected and SARS-CoV-2 infected brain assembloids, as described above, was processed on an nCounter Digital Analyzer (NanoString Technologies, Seattle, WA, USA) after hybridization to the human neuroinflammation and neuropathogenesis panel CodeSets. Three biological replicates were processed for the mock infection and each SARS-CoV-2 infection time point. Background thresholding, normalization, and t-tests were performed using nSolver Analysis Software v. 4.0.70 (NanoString Technologies Inc. Seattle, WA, USA). Background thresholding was performed using the mean of the negative controls plus two standard deviations as the threshold and setting all values below the threshold to the threshold value. Positive control normalization factors were calculated using the geometric mean of the positive controls, and any samples with a positive control normalization scaling factor greater than three-fold were excluded. The geometric mean of housekeeping genes CCDC127, CNOT10, CSNK2A2, GUSB, LARS, MTO1, SUPT7L, and TBP was used to calculate the housekeeping gene normalization factor. Any samples with a housekeeping gene normalization scaling factor greater than ten-fold were excluded. Normalized NanoString gene expressions for each infected time point were calculated using the data from the mock-infected brain assembloids as the control group. Normalized NanoString gene expression data was calculated using mean values and converted into log_2_ fold-changes. An unpaired, two-tailed t-test was performed on the log_2_ fold-change data, and changes in gene expression between mock infection and each SARS-CoV-2 infection time point were considered significant if the resulting p-value was 0.05 or less, or more preferably 0.01 or less, and there was at least a 2-fold upregulation or downregulation in the gene transcript levels. The −log_10_ p-values were calculated and plotted against the log_2_ fold-change for each gene.

### Immunocytochemistry, Imaging and Quantification

Cultured cells were washed in PBS and fixed in 4% paraformaldehyde for 15 min. Cells were blocked with blocking buffer containing 10% v/v donkey serum and 0.2% v/v Triton X-100 in PBS. Primary antibodies diluted in blocking buffer were incubated overnight at 4°C, washing three times in blocking buffer, treatment with secondary antibody (Invitrogen) were applied for 1 hr and washing three times in blocking buffer. After staining, coverslips were mounted on glass slides using prolong gold antifade reagent (Invitrogen). Primary antibodies are: Human specific Tmem119 (R&D Systems, MAB10313), Human specific Iba1 (R&D Systems, MAB7308). The following cyanine 2 (Cy2)-conjugated secondary antibodies were used to detect the first antibodies: donkey antibody against mouse (Invitrogen).

### RNA Seq

Total RNA was harvested from culture using Trizol (Invitrogen) following the manufacturer’s protocol. Ribosomal RNAs were subtracted from total RNA using RiboZero magnetic kit (Epicentre). Libraries from rRNA-subtracted RNA were generated using ScriptSeq 2 (Epicentre) and sequenced on an Illumina HiSeq 2500. Reads were mapped to the human genome (release 19) using TopHat2. Reads were annotated using a custom R script using the UCSC known Genes table as a reference. Gene ontology shows output from a enrichment analysis on DAVID (david.abcc.ncifcrf.gov/), searching the following categories: GO Biological Process, GO Molecular Function, PANTHER Biological Process, PANTHER Molecular Function, KEGG and PANTHER pathway. Enrichment terms and associated genes and statistics are presenting with additional statistical analyses also provided.

### Deconvolution of Bulk RNA-seq Using Single-cell Reference

To estimate the relative cell-type composition of immune-vascularized forebrain organoids (FORMA-COs), we applied a reference-based deconvolution approach using CIBERSORTx (https://cibersortx.stanford.edu), a computational tool that infers cell-type proportions from bulk RNA-seq data using single-cell transcriptomic references.

We constructed a reference signature matrix from the publicly available single-cell RNA-seq dataset reported by Trevino et al. (*Cell*, 2021; PMID: 34390642), which profiles chromatin and gene-regulatory dynamics of the developing human cerebral cortex. This dataset includes single-cell transcriptomes from midgestation human cortical tissue spanning multiple neural, glial, and vascular cell types. Raw counts were downloaded from the Gene Expression Omnibus (GEO accession: GSE162170) and processed using Seurat (v4.3.0). Cells were filtered for quality control, normalized using SCTransform, and annotated according to cell-type labels provided in the original publication (e.g., radial glia, intermediate progenitors, excitatory neurons, interneurons, astrocytes, oligodendrocyte lineage, microglia, endothelial cells, and pericytes). To generate the CIBERSORTx signature matrix, we selected the top 50 marker genes per cell type based on average log fold-change and adjusted p-value (Wilcoxon rank-sum test) and input this matrix into CIBERSORTx’s “Create Signature Matrix” module. The resulting signature was used in the “Impute Cell Fractions” module to deconvolute bulk RNA-seq profiles of APOE isogenic FORMA-COs. Bulk RNA-seq data were normalized to transcripts per million (TPM) prior to analysis, and batch correction mode (S-mode) was enabled to account for platform differences between single-cell and bulk datasets. Visualization of inferred compositions was performed using bar plots and PCA clustering in R (v4.2.2). The robustness of cell-type estimations was confirmed by cross-validating with immunostaining and transcriptomic marker gene expression in matched organoid samples.

### Statistics

All quantitative data are presented as mean ± SEM. “n” refers to the number of independent biological replicates (separate iPSC differentiations or individual animals), unless otherwise stated. Statistical analyses were performed using GraphPad Prism v9.0 (GraphPad Software) or R v4.2.0. Normality of each dataset was assessed by the Shapiro–Wilk test. For comparisons between two groups, unpaired two-tailed Student’s t-tests were used when data were normally distributed; otherwise, the nonparametric Mann–Whitney U test was applied. For multiple group comparisons, one-way or two-way ANOVAs were performed as appropriate, followed by Tukey’s or Šidák’s post hoc tests to correct for multiple comparisons. When comparing viral replication kinetics or time course measurements, repeated-measures two-way ANOVA was employed with Geisser–Greenhouse correction. Categorical data were analyzed by chi-square or Fisher’s exact tests, as indicated. Correlations were assessed by Pearson’s or Spearman’s rank correlation coefficients based on data distribution. A p-value < 0.05 was considered statistically significant in all analyses. Exact p-values and details of each test are reported in the figure legends. Wherever feasible, experiments were randomized and investigators blinded to genotype or treatment during data collection and analysis to minimize bias.

## Acknowledgment

We gratefully acknowledge the generous support of the National Institutes of Health (R37 NS067525, AG066507, R01 AI161829-A1, R01 AI145435-A1) and Emergent Ventures at the Mercatus Center, George Mason University (Grant #2167).

---

**Figure S1.**
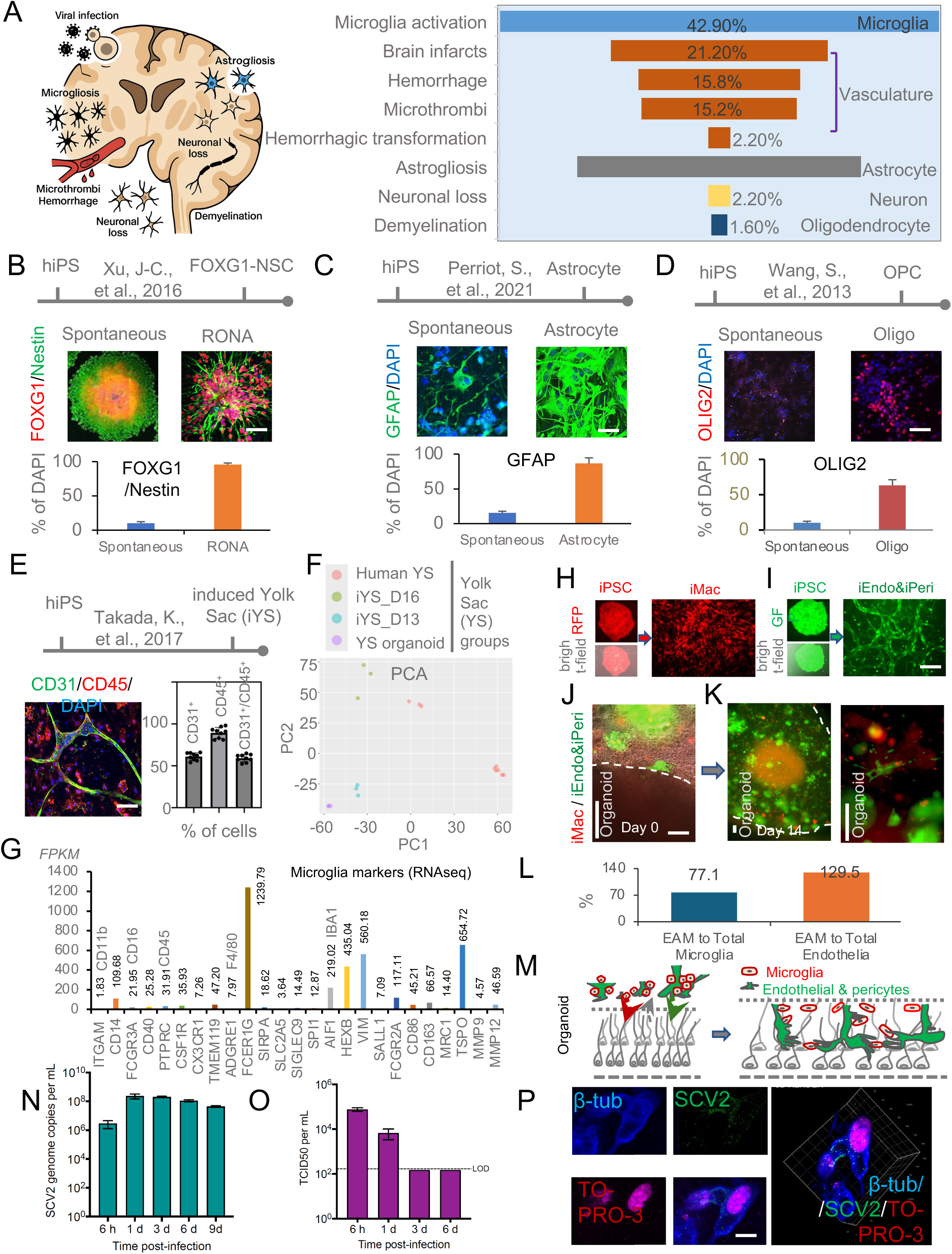
**Characterization of FORMA-COs and viral susceptibility.** (A) Summary of human brain IHC showing SCoV2, vascular injury, and gliosis. (B–D) FOXG1⁺/Nestin⁺ neural stem cells show multipotency (GFAP⁺, Olig2⁺). (E–G) Yolk sac-like progenitors express CD31, CD45, and microglial markers. (H–K) Fluorescent tracking of iMac and iEndo/iPeri integration. (L–M) Schematic and quantification of endothelial-associated microglia. (N–O) Viral RNA and titer dynamics post-infection. (P) Confocal imaging shows SCoV2 colocalization with neurons.

**Figure S2.**
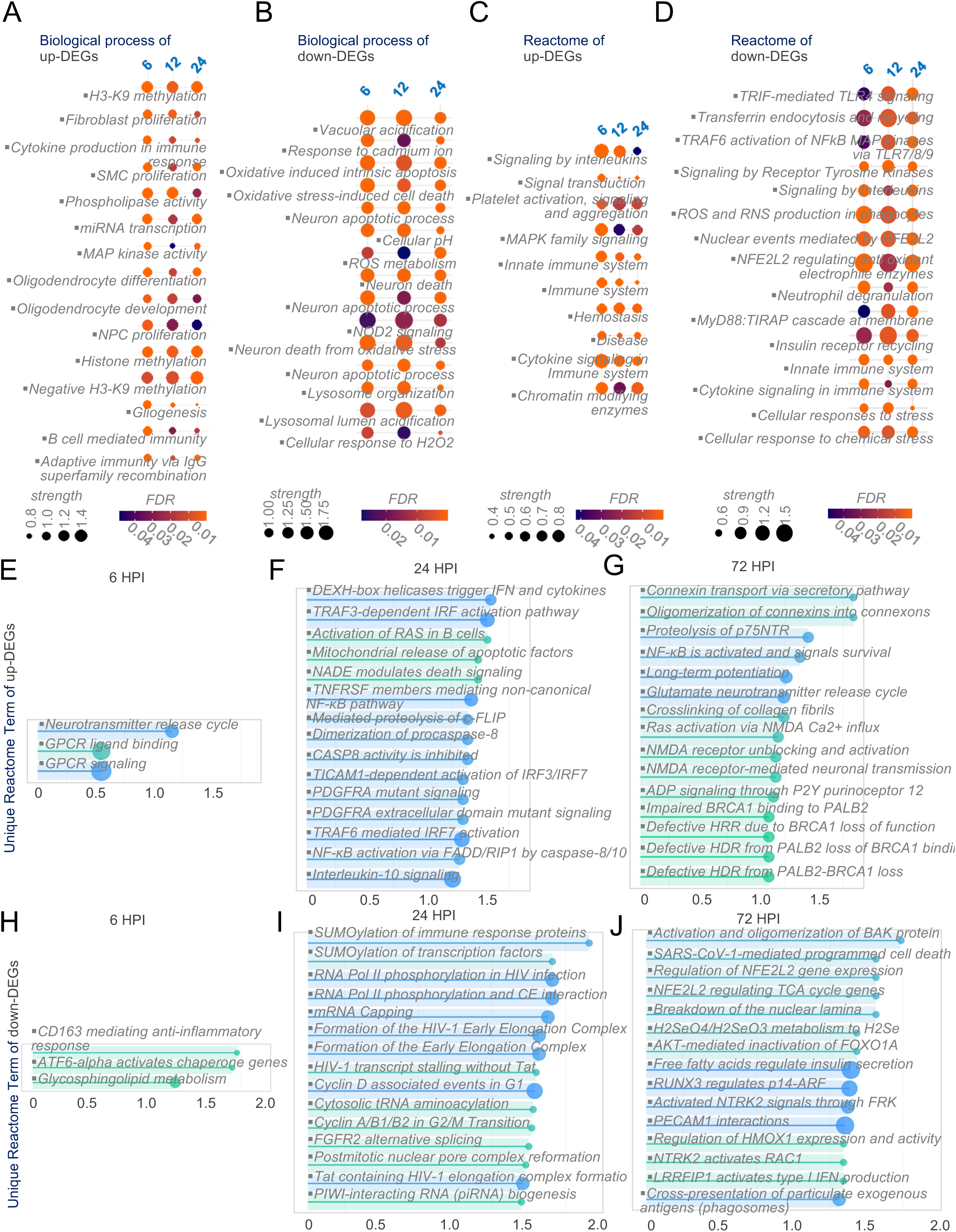
**Pathway enrichment in SCoV2-infected FORMA-COs.** (A–B) GO analysis shows time-dependent immune activation, gliogenesis, oxidative stress, and apoptosis. Early response includes innate immunity; later stages involve remodeling and neuroinflammation. (C–D) Reactome pathways confirm early cytokine signaling and persistent innate immune activation. (E–F) At 6 and 24 HPI, interferon signaling, apoptosis, and NF-κB activation dominate; downregulated genes involve anti-inflammatory responses and transcriptional repression. (G–J) At 72 HPI, enriched pathways include disrupted synaptic and gap junction signaling, impaired DNA repair, and vascular dysfunction. Dot size indicates gene count; color reflects FDR; x-axis shows enrichment strength.

**Figure S3.**
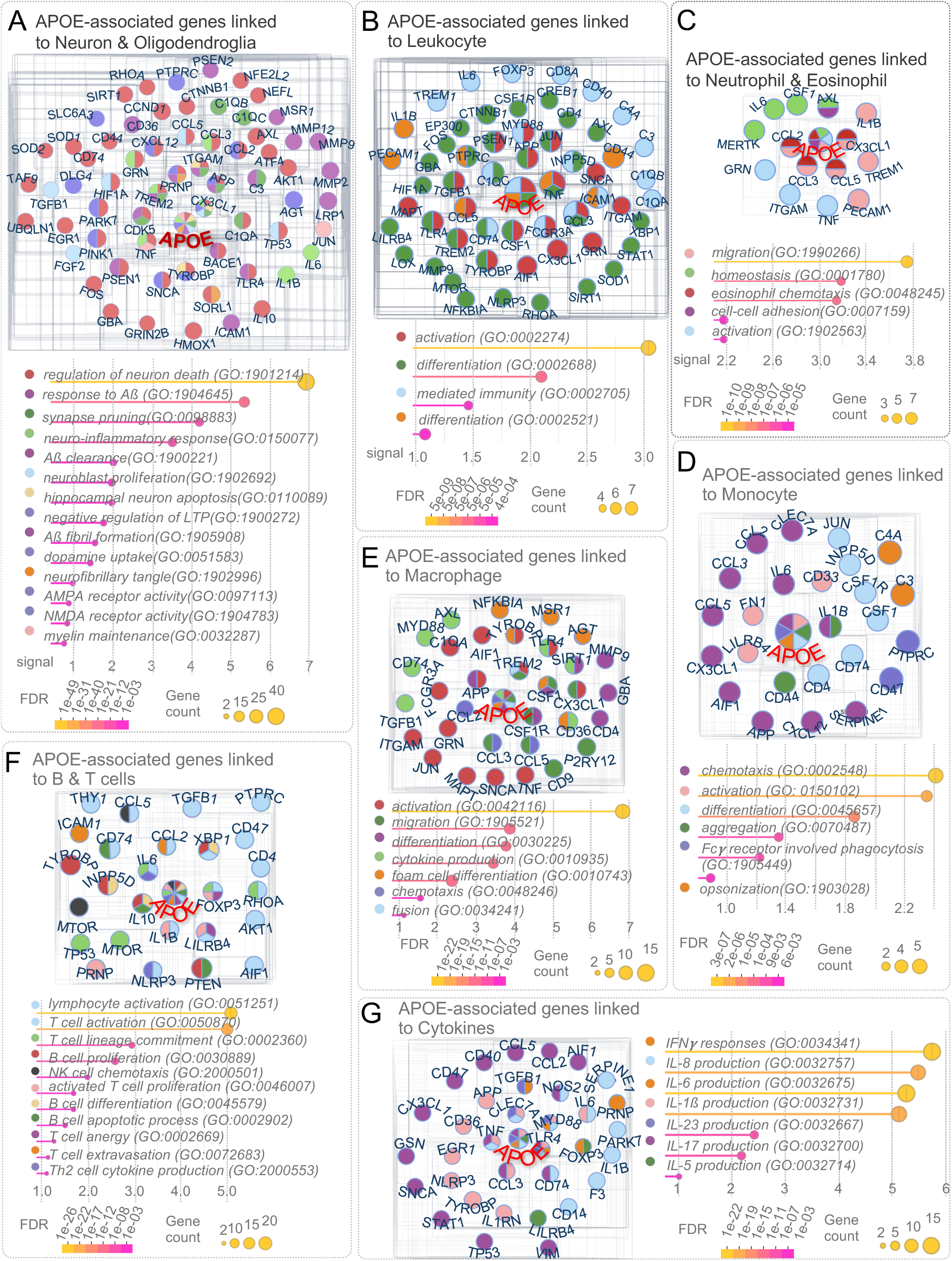
**Functional segregation of APOE interactomes across CNS and immune cells.** (E) APOE-centered network enriched for synaptic, myelination, and glial function. (F) Leukocyte-specific APOE network highlights activation, migration, and adhesion pathways. (C–E) APOE modulates granulocyte, monocyte, and macrophage programs including chemotaxis, cytokine signaling, and phagocytosis. (F) Adaptive immunity network shows APOE links to B and T cell activation and immune regulation. (G) Cytokine-centered network shows enrichment for interferon and interleukin responses. Pie charts indicate multifunctional gene roles; dot plots reflect GO term enrichment by gene count and FDR.

**Figure S4.**
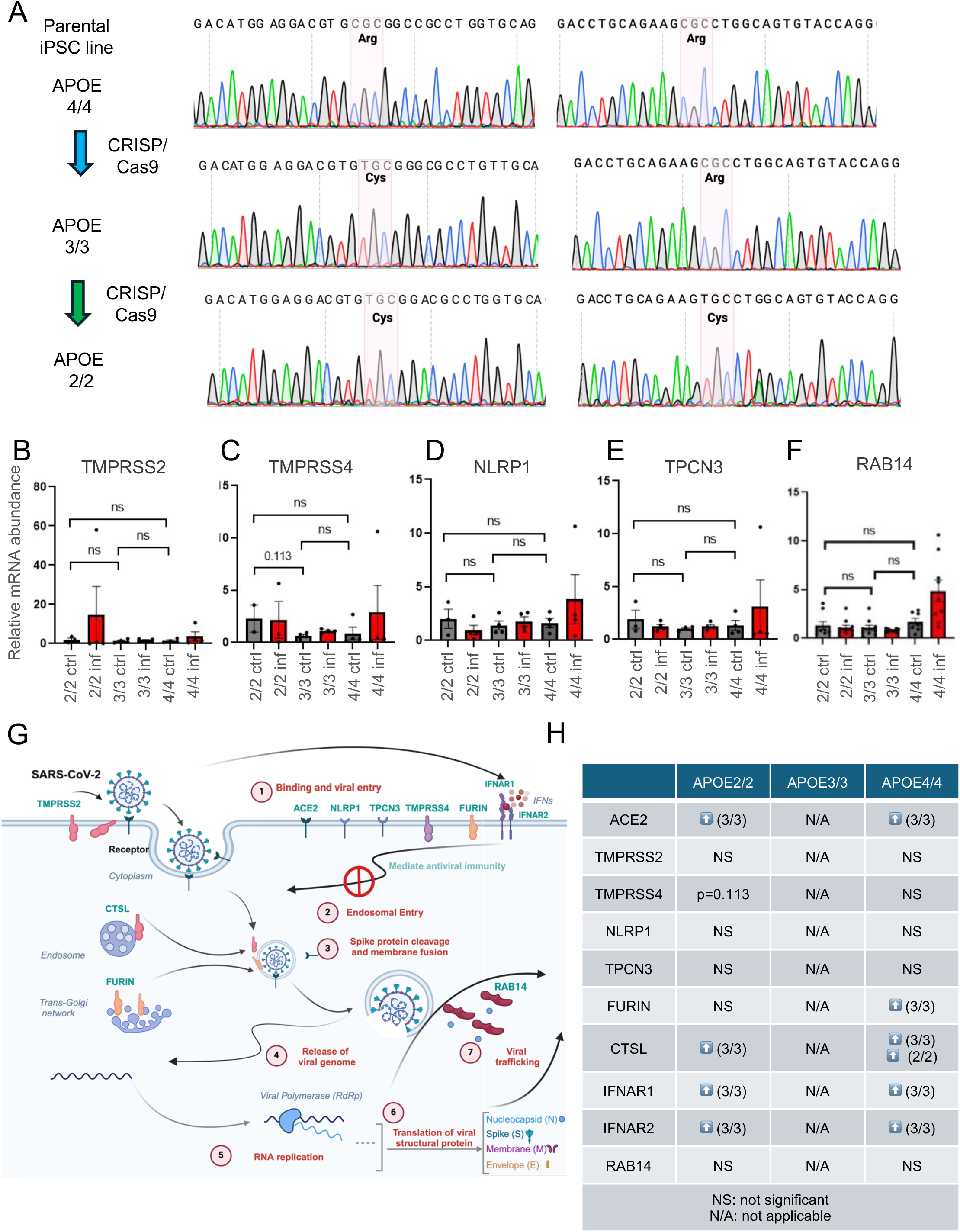
**APOE Genotype Regulates Host Viral Entry and Immune Response Genes.** (A) Sanger sequencing confirms APOE genotypes via rs429358 and rs7412 SNPs. (B-F) Bar graphs show expression of TMPRSS2, TMPRSS4, NLRP1, TPCN3, and RAB14 across APOE genotypes under control/infected conditions. (G) Schematic of SARS-CoV-2 life cycle and host entry factors. (H) Summary table of APOE-dependent expression of viral entry and immune genes. APOE4/4 and 2/2 cells upregulate ACE2, CTSL, IFNAR1/2, and FURIN compared to APOE3/3; TMPRSS2 and others show no genotype effect.

**Figure S5.**
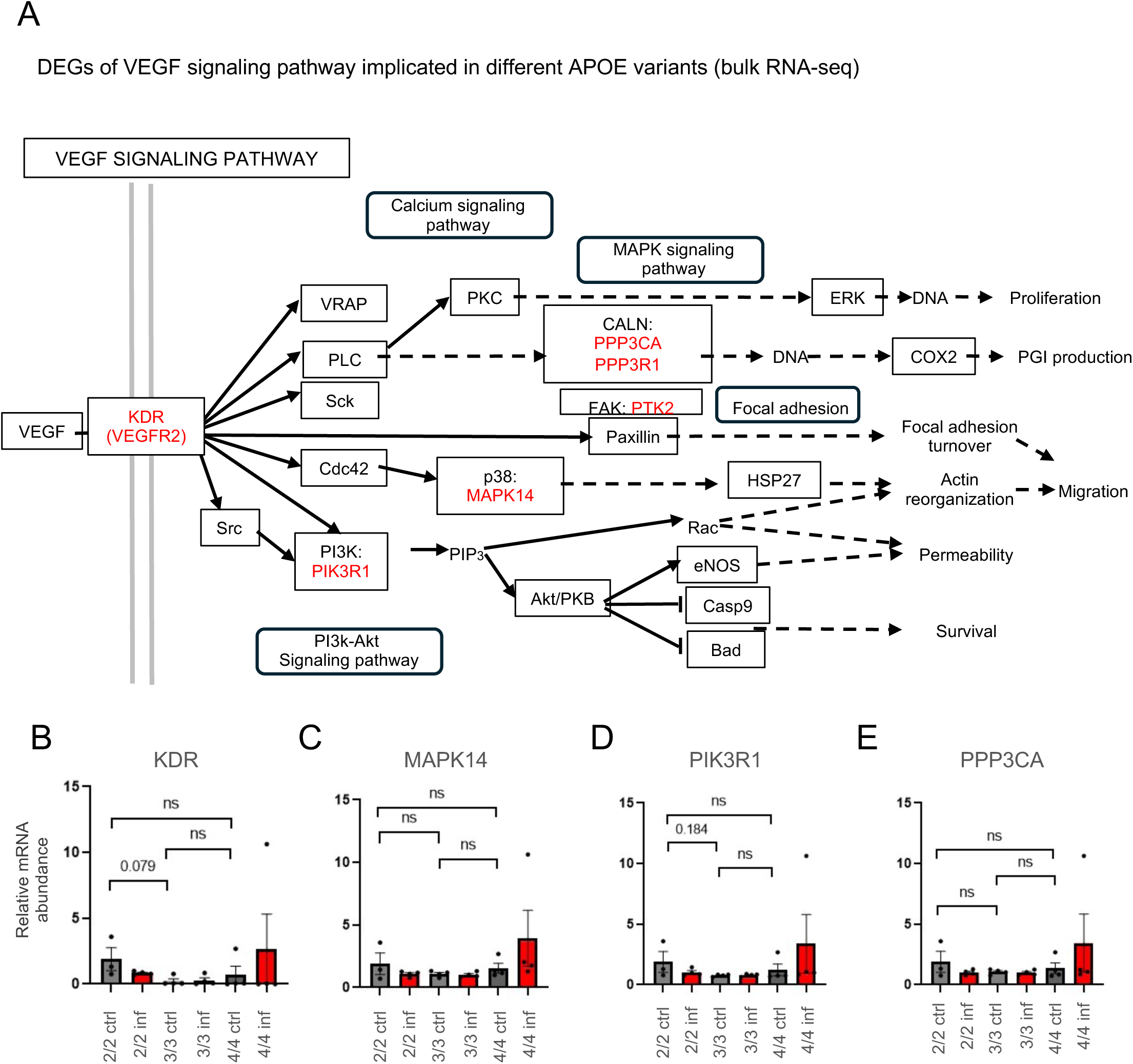
**APOE Genotype Alters VEGF Pathway Components Under Infection.** (B) Schematic of VEGF signaling with genotype-modulated components (purple). Differential expression seen in VEGFA, PTK2, and PPP3R1. (C) KDR (VEGFR2) mRNA shows a trend toward increase in APOE2/2 controls (p = 0.079) and slight rise in APOE4/4-infected cells. (D) MAPK14 (p38) shows modest, nonsignificant elevation in APOE4/4-infected group. (E) PIK3R1 shows trending increases in APOE4/4-infected and APOE2/2 controls (p = 0.184). (F) PPP3CA levels remain stable; slight increase in APOE4/4-infected cells is not significant. Trends suggest genotype-related activation potential for VEGF downstream pathways.

**Figure S6.**
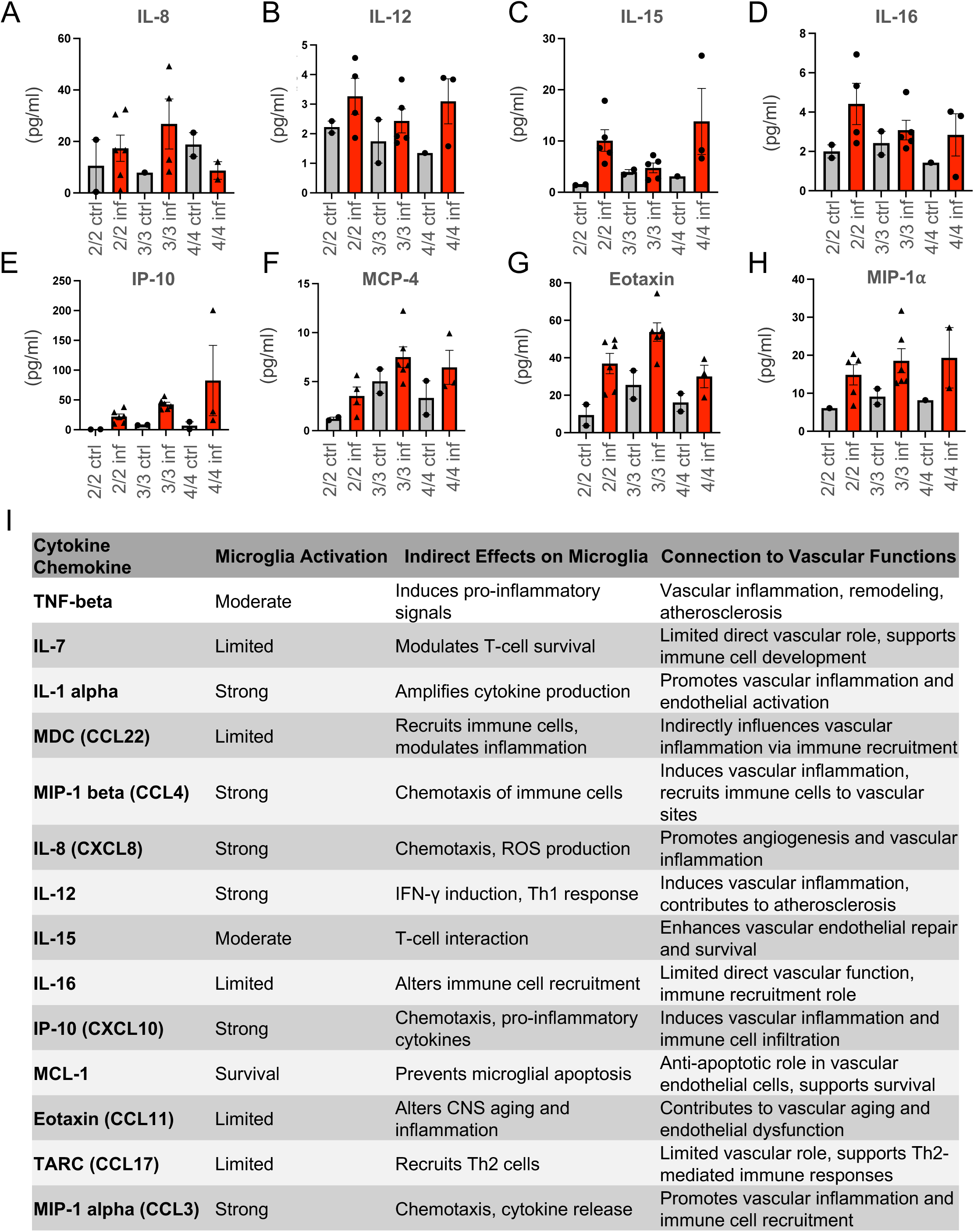
**APOE Genotype Differentially Modulates Cytokine and Chemokine Responses.** (A–B) IL-8 and IL-12 levels rise with infection across genotypes; IL-8 highest in APOE3/3; IL-12 elevated in APOE4/4. (C–D) IL-15 and IL-16 increase in APOE4/4 and APOE2/2 post-infection; IL-16 highest in APOE2/2. (E–F) IP-10 and MCP-4 show increased levels in APOE4/4 and APOE2/2. (G) Eotaxin highest in APOE3/3, lower in APOE2/2. (H) MIP-1α increases in all genotypes; APOE4/4 shows strongest induction. (I) Summary chart categorizing cytokines by microglial impact, immune recruitment, and vascular relevance.

## Notes

### Competing Interest Statement

The authors have declared no competing interest.

